# Modeling Host-Virus Interactions of Avian Corona Virus Reveals Cytokine Feedback as a driver of Strain-Specific Immune Response

**DOI:** 10.64898/2026.11.20.768156

**Authors:** Krishanu Mondal, Motamed Elsayed Mahmoud, Tushar Pal, Santidan Biswas, Dipjyoti Das, Mohamed Faizal Abdul-Careem

## Abstract

The interplay between RNA viral dynamics and innate immunity remains poorly understood, mainly because RNA viruses trigger highly nonlinear cytokine responses. We address this using infectious bronchitis virus (IBV), a major poultry pathogen, as a model system. We developed a set of mathematical models and compared those with experimental data from two IBV strains infecting chicken macrophages. By model fitting and supervised Machine Learning we reveal that positive feedback in interferon-*α* production is essential to reproduce the dynamics of one IBV strain but likely absent in the other. We further provide strain-specific estimations of key parameters such as infection rates, production and decay rates of virus and cytokine levels. Notably, different strains produced distinctive cytokine profiles, highlighting feedback as a key determinant of strain-specific immune signatures. Our results establish a quantitative framework for dissecting host-virus interactions and uncover cytokine feedback as a generic yet underappreciated feature in models of RNA virus kinetics.

## Introduction

Mathematical modeling provides a rigorous framework to understand the process of viral infection and replication within a host. Over the past few decades, numerous models of viral dynamics have been developed to capture the within-host behavior of both RNA and DNA virus (1–5). Viral dynamic models have been extensively applied to infections such as Influenza virus (1, 6), HIV (7) , Hepatitis virus (8, 9), Zika virus (10) , and more recently, SARS-CoV and SARS-CoV-2 (11–13). Recent efforts (14) have demonstrated the application of such models in understanding the early-stage dynamics of SARS-CoV-2 infection. Collectively, these models not only provide insights into the within-host evolution of viral infections but also inform clinical decision-making and public health strategies through simulation of therapeutic interventions and vaccination scenarios.

The classical framework used to capture the dynamics of viral infections, both *in vivo* and *in vitro*, is the three variable target cell limited model (1, 3, 15). This model describes the interactions between susceptible target cells (T), which become infected at a certain rate upon exposure to free virions (V), and the newly infected cells (I) that start producing new virus particles. This model explains certain aspects of many viral infections in a coarse-grained way and provide partial insights; however, it does not consider the complex interplay between viral production and the innate immune system. In response to the entry of a foreign pathogen, the host immune system elevates cytokines such as interferons (IFN), tumor necrosis factors (TNF), and multiple interleukins (ILs), which play critical roles in initiating and regulating immune responses (16, 17). Recognizing the critical immunomodulatory role of cytokines, subsequent models explicitly included interferons (IFN) to offer a more complete representation of host–virus interactions (18–20). For example, a recent model proposed a delayed cytokine production that successfully reproduced the experimentally observed bimodal peaks in the viral titer versus time curves. Some recent models have also incorporated the kinetics of adaptive immune response (21, 22). However, a key limitation in most subsequent works is the absence of simultaneous fitting to both viral concentration and cytokine expression data. Moreover, these models often omit the feedback control mechanisms that are known to influence interferon (IFN) production (23, 24). Despite the inclusion of such feedback in some modeling studies (25), the lack of fitting to experimental datasets limited the model to theoretical exploration only.

Here, we study avian coronavirus, infectious bronchitis virus (IBV) as a model to examine the interplay between viral infection and the innate immune system. Infectious bronchitis (IB) caused by IBV is a major respiratory disease of chickens that can also affect the urinary and reproductive tracts, leading to substantial production losses in commercial broiler and layer flocks worldwide (26–28). The morbidity rates due to IBV can reach up to 100% in young birds (2–3 weeks of age) (29). The causative agent, IBV, belongs to the family *Coronaviridae*. Owing to it’s RNA genome and high rates of mutation and recombination, IBV exists as multiple serotypes and strains (30).

Despite considerable progress in viral dynamic models for human pathogens, the mathematical modelling of viral infections in animals remains relatively underexplored. Only a small number of research have focused on animal hosts, including horses (6), avian influenza in poultry (31–33), livestock (34), and bats (35). Animal infections often hold zoonotic potential, making it vital to understand the viral dynamics inside animal hosts. As many emerging human viruses originate from animal reservoirs (36–38) systematic studies of animal viral infections, particularly in poultry animals, are crucial for identifying fundamental mechanisms governing viral replication, immune regulation, and cross-species transmission. In this regard, IBV provides an important avian model for quantitatively exploring host–virus interactions.

To investigate the within-host kinetics of IBV in avian systems, we developed mechanistic mathematical models and validated with experimental data obtained from IBV-infected chicken macrophage cell lines. We developed two models where we explicitly included IFN kinetics and feedback in type I IFN production. By systematically comparing these model formulations through parameter estimation and data fitting, we tested the hypothesis of whether feedback control in IFN production is necessary or not for an efficient clearance of viral infection. We tested our modelling framework using two IBV strains with distinct pathogenic characteristics: the classical respiratory strain Connecticut A5968 (Conn A5968) and the nephropathogenic strain Delmarva/1639 (DMV/1639) (39, 40). Through model fitting and supervised Machine Learning, we conclude that positive feedback in type I IFN production is essential to reproduce the viral dynamics of one strain (DMV/1639) but likely absent in another strain (Conn A5968). Thus, cytokine-dependent feedback affects the immune response in a strain-specific way. Additionally, different virus strains generate quantitatively distinct cytokine profiles, underlining cytokine feedback as a fundamental driver of strain-specific immunological response. Together, our findings provide a quantitative framework for analyzing host–virus interactions and highlight cytokine feedback as a broadly relevant yet inadequately investigated component of RNA virus dynamics.

## Results

### A virus–target cell–interferon model captures the viral dynamics for one strain but not for the other

We first developed a mathematical model to describe the dynamics of IBV infection in a chicken macrophage cell line. We assume that the immune system modulates viral dynamics by suppressing the viral replication via interferon signaling. To incorporate this process, we formulated the following Model 1 (Fig. 1A) that represent the interactions among target cells (*T*), extracellular virus (*V*), and IFN expression (*F*). This Model 1can be described by the following equations

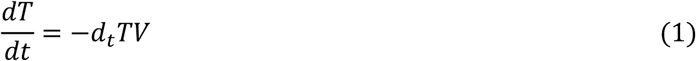

**Figure 1.**
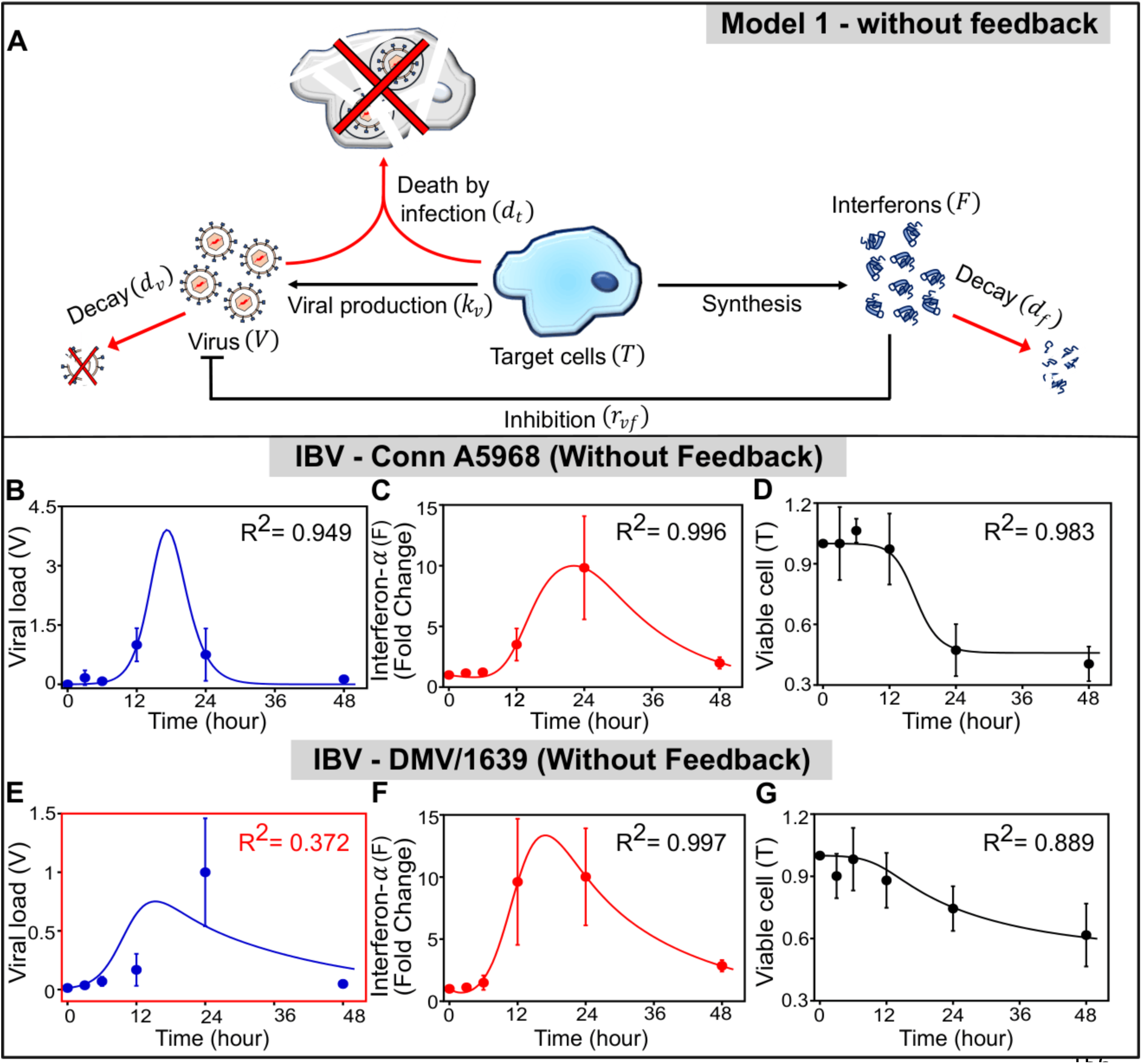
A three-variable model can predict the viral dynamics of one strain but cannot explain the dynamics of another strain. **(A)** A schematic representation of our three-variable model (Model 1) describing the viral dynamics coupled with the innate immune response. The free virus (V) infects target cells (T), which then produce virus as well as interferons (F) that further inhibit viral production (see Equations 1-3). Arrows indicate activation or production processes, while blunt-ended lines denote inhibitory interactions. **(B–G)** Temporal dynamics of viral load, interferon-α, and viable cells are plotted for the IBV-Conn A5968 (B-D) and IBV-DMV/1639 (E-G) strains. Solid lines represent model predictions, and filled circles indicate experimental measurements (n = 3; error bars indicate standard deviation). The coefficients of determination (*R*^2^), indicating the goodness of fit, are shown in the right upper corner of each panel. Panel E for IBV-DMV/1639, highlighted in a red box, shows a poor fit to the model prediction, with a much lower *R*^2^ value.

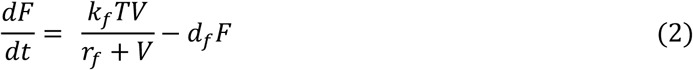

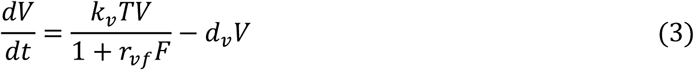

Here, the parameter *d_t_* represents the rate of infection when the virus encounters susceptible target cells; *d_f_* and *d_v_* represent the decay rates of IFN and virus, respectively; while *k_f_* and *k_v_* are the production rates of IFN and virus, respectively. Interferons inhibit the virus production, with *r_vf_* representing the strength of this inhibition.

To compare model predictions with experimental data, we consistently scaled both datasets. Specifically, the extracellular IBV RNA copy number (a measure of the viral load) was normalized by the experimentally measured maximum mean value near the peak, the viable cell fraction was normalized by its initial value, and extracellular IFN by its initial value in the model and by its expression level in mock-infected (MOCK) cultures in the experiments (see Materials and Methods in SI for details). We estimated the model parameters by fitting model predictions to the experimental time-course data for extracellular viral load, interferon (IFN) fold change, and the proportion of viable target cells (see Materials and Methods in SI). The best-fit parameter values (Table 1) were obtained assuming that, at an experimental multiplicity of infection (MOI) of 0.1, 50% of the initial viral particles were in the extracellular space, while the remaining 50% initiated infection at the intracellular level. We also verified that varying the initial extracellular virus fraction did not alter the qualitative predictions of Model 1 and reproduced the experimental trend (Fig. S1).

**Table 1.**
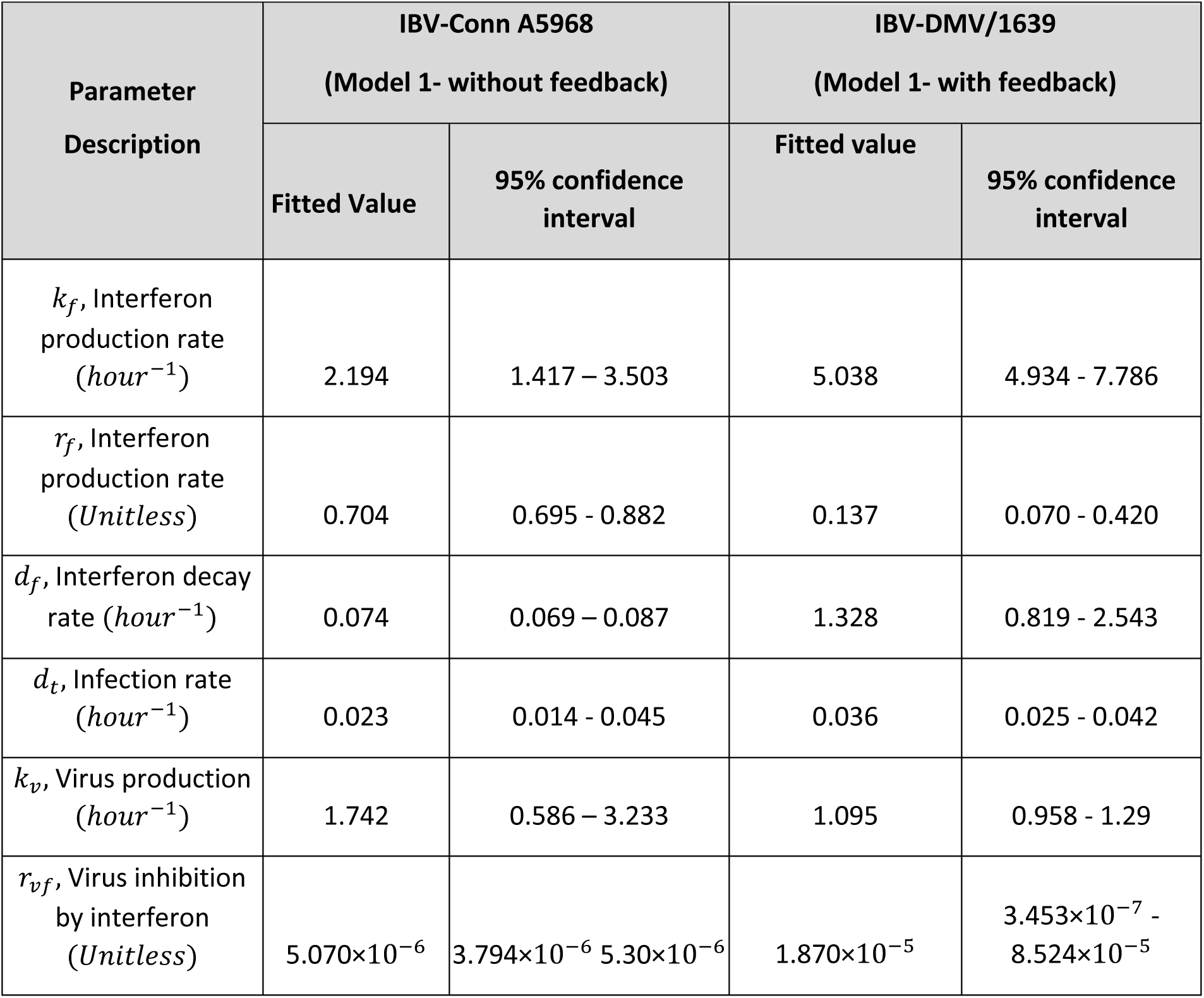
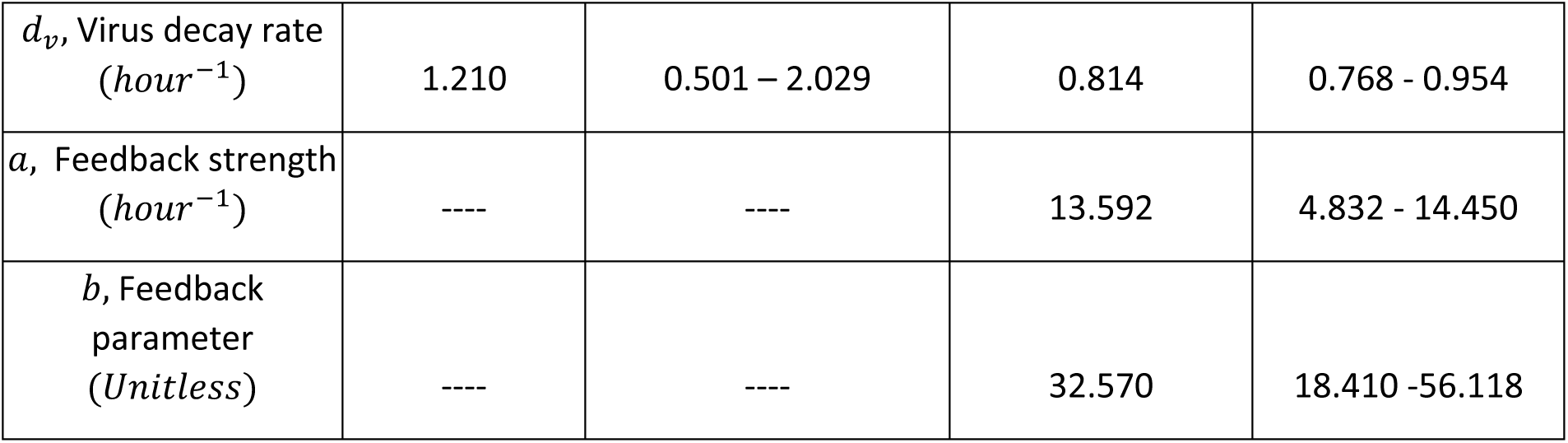
Parameter values estimated from least squares fitting for the IBV-Conn A5968 and IBV-DMV/1639 strain using Model 1 (see SI for details). Units and values are presented along with the 95% confidence intervals.

| Parameter Description | IBV-Conn A5968<br>(Model 1- without feedback) |  | IBV-DMV/1639<br>(Model 1- with feedback) |  |
|---|---|---|---|---|
|  | Fitted Value | 95% confidence interval | Fitted value | 95% confidence interval |
| *k<sub>f</sub>*, Interferon production rate (*hour*<sup>−1</sup>) | 2.194 | 1.417 – 3.503 | 5.038 | 4.934 - 7.786 |
| *r<sub>f</sub>*, Interferon production rate (*Unitless*) | 0.704 | 0.695 - 0.882 | 0.137 | 0.070 - 0.420 |
| *d<sub>f</sub>*, Interferon decay rate (*hour*<sup>−1</sup>) | 0.074 | 0.069 – 0.087 | 1.328 | 0.819 - 2.543 |
| *d<sub>t</sub>*, Infection rate (*hour*<sup>−1</sup>) | 0.023 | 0.014 - 0.045 | 0.036 | 0.025 - 0.042 |
| *k<sub>v</sub>*, Virus production (*hour*<sup>−1</sup>) | 1.742 | 0.586 – 3.233 | 1.095 | 0.958 - 1.29 |
| *r<sub>vf</sub>*, Virus inhibition by interferon (*Unitless*) | 5.070×10<sup>−6</sup> | 3.794×10<sup>−6</sup> 5.30×10<sup>−6</sup> | 1.870×10<sup>−5</sup> | 3.453×10<sup>−7</sup> - 8.524×10<sup>−5</sup> |
| *d<sub>v</sub>*, Virus decay rate (*hour*<sup>−1</sup>) | 1.210 | 0.501 – 2.029 | 0.814 | 0.768 - 0.954 |
| *a*, Feedback strength (*hour*<sup>−1</sup>) | ---- | ---- | 13.592 | 4.832 - 14.450 |
| *b*, Feedback parameter (*Unitless*) | ---- | ---- | 32.570 | 18.410 -56.118 |

As shown in Figs. 1B–1D Model 1 successfully captured the observed dynamics of viral load, IFN response, and target-cell viability for the IBV-Conn A5968 strain (with R² values exceeding 0.7). However, Model 1 could not simultaneously reproduce all three experimental observables for the IBV-DMV/1639 strain (Figs. 1E–1G). Although Model 1 reasonably captured the IFN response and viable-cell fraction (Figs. 1F and 1G), it’s fit to the viral load data remained poor (with R² ∼ 0.4; Fig. 1E).

### Inclusion of feedback on cytokine production is necessary to explain the strain-specific viral dynamics

The inability of Model 1 to simultaneously reproduce all experimental observations for the IBV-DMV strain suggests that important biological processes were not adequately represented in the original framework. To overcome this limitation, we extended Model 1 to incorporate positive feedback in cytokine production (Fig. 2A), replacing Eq. 2 with the following equation:

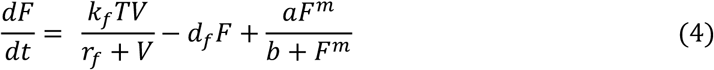

**Figure 2:**
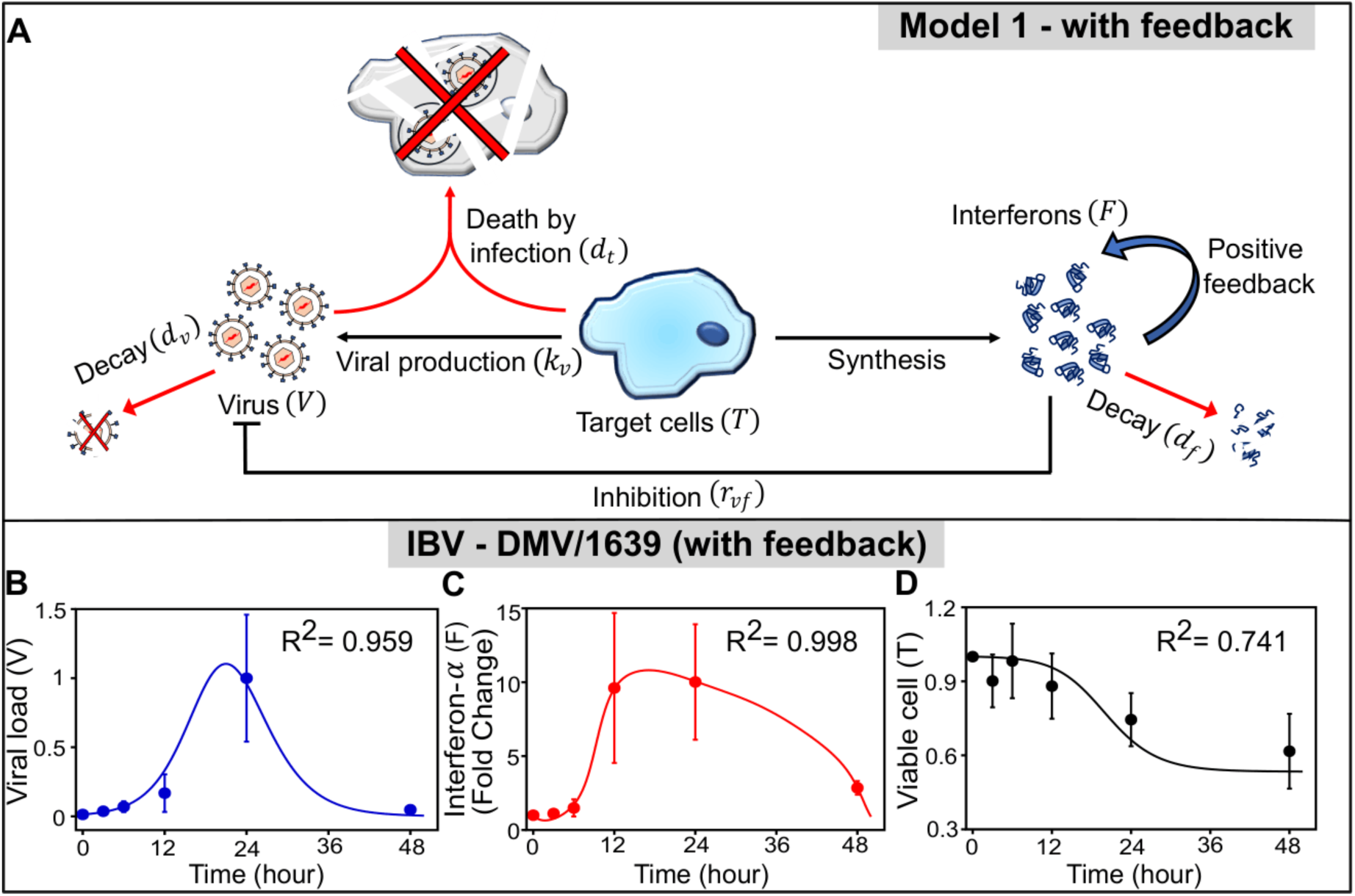
Inclusion of positive feedback to interferon production in Model 1 captures the infection dynamics of the IBV-DMV strain. (A) A schematic of modified Model 1 with a positive feedback loop in the interferon dynamics, modeling the stimulatory effects of interferon on its own production (see Equation 4). (B–D) Comparison of model predictions and experimental data for the IBV-DMV/1639 strain with the positive feedback. Panels show the temporal dynamics of (B) viral load, (C) interferon-α concentration, and (D) viable cell count over a 48-hour period post-infection. Solid black lines represent Model solutions, and filled circles indicate experimental data (n = 3 per time point). Error bars represent standard deviations. *R*^2^ values are shown in the top corner of each panel.

In Equation (4), the positive feedback in interferon is represented by a Hill-like function, where, *a* and *b* are the feedback parameters, and *m* is the Hill coefficient. Throughout this study, we primarily set *m* = 2, as the case ‘*m*=1’ failed to reproduce the IBV-DMV/1639 data (Fig. S2A–S2C). Subsequently, we also explored the possibility of inferring (m) from the data by varying it between 1 and 3 in a supervised machine learning (ML) framework.

The modified Model 1 was then solved numerically for the IBV strain using Eqs. 1, 3, and 4. Incorporating positive feedback in cytokine production substantially improved the agreement between model predictions and experimental observations for the IBV-DMV/1639 strain (Fig. 2B–2D). We also assessed the robustness of this result to the assumed initial distribution of viral particles by varying the fraction of initial extracellular viral load. The modified model produced satisfactory agreement with the experimental data (Fig. S3).

The best-fit parameter estimates for the IBV-Conn A5968 strain, obtained using the minimal Model 1, are reported in Table 1; while the optimized parameters for the IBV-DMV/1639 strain were obtained using the modified Model 1 with feedback. Notably, the estimated feedback parameter (*b*) for IBV-Conn A5968 was nearly zero (*b*∼10^−9^; see Eq. 4), suggesting that positive feedback in cytokine production is negligible for this strain (Fig. S2D-S2I).

### A modified target-infected cell model can describe the viral dynamics of one specific strain in the absence of cytokine feedback

Despite providing a satisfactory overall fit to the experimental data, Model 1 did not accurately capture the viral-load dynamics near the peak for the IBV-Conn A5968 strain (Fig. 1B). Experimentally, the viral loads at 12 and 24 h post-infection were comparable, indicating a plateau-like peak between these time points. In contrast, Model 1 predicted a sharper peak, potentially overestimating viral load during this peak phase. This discrepancy suggests that the model may not fully capture the biological processes regulating viral accumulation, especially near the peak of infection.

To address this limitation, we extended the framework to develop Model 2 (Fig. 3A), following a previous study (6). Model 2 explicitly describes the transition of macrophages from an uninfected state to a virus-producing infected state. We assume that free extracellular virions infect susceptible, uninfected cells (*U*), converting them into infected cells (*I*), which subsequently serve as the primary source of viral production. Eqs. 5–8 below describe the dynamics of Model 2.

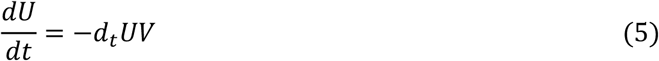

**Figure 3:**
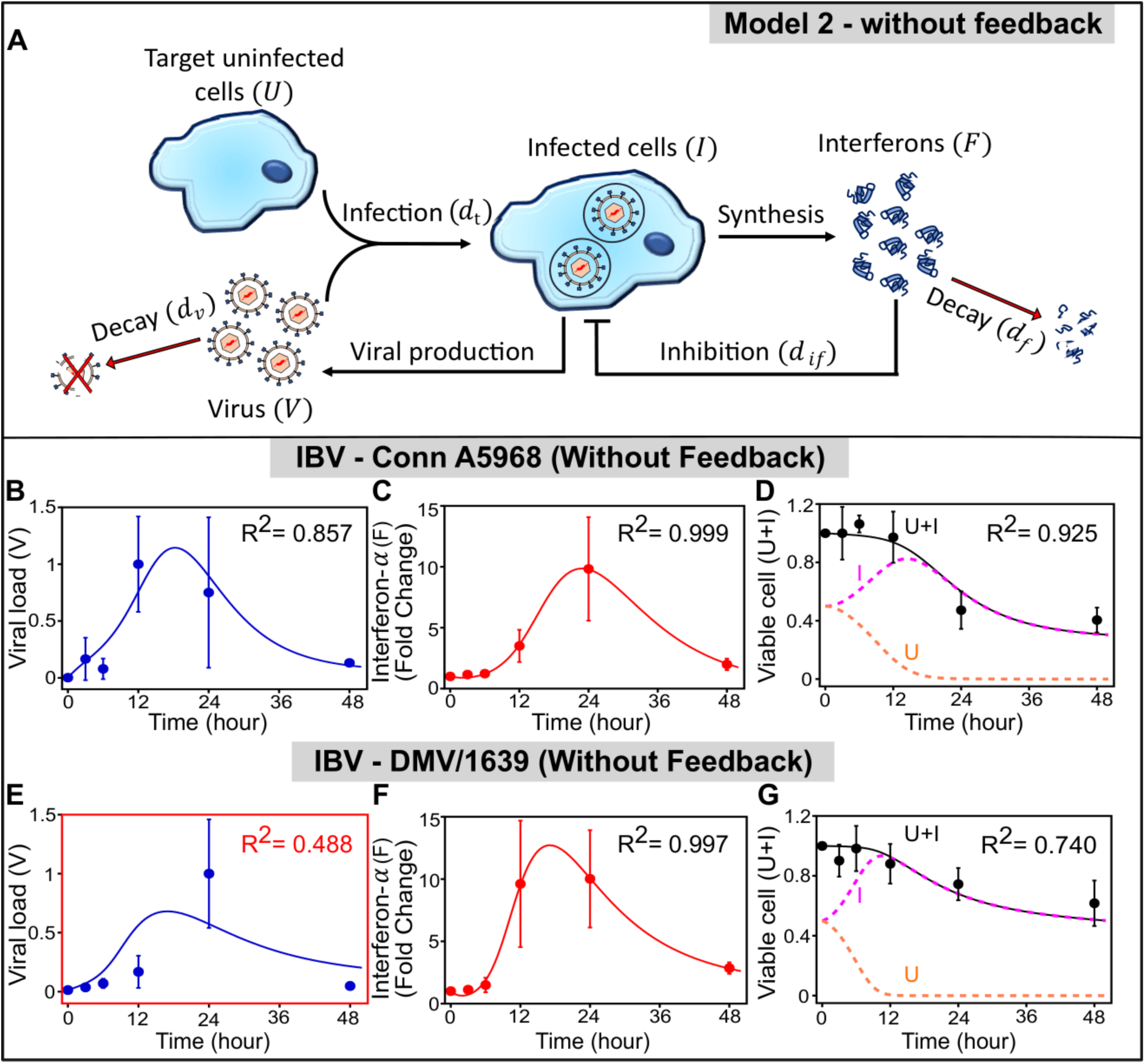
Model 2 with infected and uninfected cell populations can describe the infection dynamics of the IBV-Conn strain, but cannot reproduce the IBV-DMV dynamics. **(A)** The schematic representation of our four-variable model (Model 2), incorporating more biological details. Here, the free virus (V) infects susceptible uninfected cells (U), resulting in infected cells (I). Infected cells then produce both virus and interferons (F), which induce an antiviral effect by inhibiting the production of infected cells (see Equations 5-8). Arrows denote production or activation, while blunt-ended lines indicate inhibitory effects. **(B-G)** Comparison of model predictions and experimental data for the IBV-Conn A5968 (B-D) and IBV-DMV/1639 (E-G) strains. Solid lines represent model predictions, and filled circles indicate experimental measurements (n = 3 per time point; error bars indicate standard deviation). The magenta and orange dashed curves in panels D and G represent the predicted numbers of infected and uninfected cells, whose sum equals the total viable cells, as compared with the data. Panel E for IBV-DMV/1639, within a red box, highlights a poor fit with a relatively low coefficient of determination.

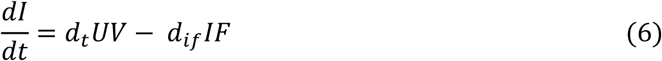

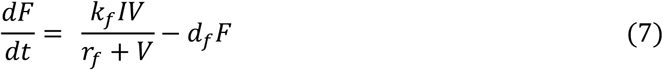

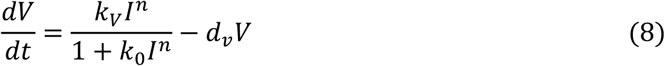

Here, *d_t_* is the transition rate from uninfected cells (*U*) to infected cells (*I*). Explicitly distinguishing these two cell populations makes the model more biologically realistic, as it brings an implicit delay in the viral production after initial infection, a general feature of the viral life cycle (41). The parameter *d_if_* denotes the clearance rate of infected cells mediated by interferon. Here we modelled the growth term of the virus by a hill function dependent on infected cells, where *n* is the Hill coefficient, and *k_V_*, *k*_(_ are the scaling and saturation parameters, respectively. Throughout this study, we fixed *n* = 3.

To examine the effect of the implicit delay in viral production following infection, we numerically solved Model 2, assuming equal initial proportions of uninfected and infected cells (0.5 each). This proportion, corresponding to the initial ratio of extracellular-to-intracellular viral distribution was held fixed throughout the study (see Methods in SI). However, varying the initial population proportions yielded similar predictions and fits with data, indicating that our results were robust (Fig. S4 and S5).

Figure 3 shows the best-fit predictions of Model 2. Similar to Model 1, Model 2 successfully captured the time-series data of viral load, IFN fold change, and viable cell proportion for the IBV-Conn A5968 strain (Fig. 3B–3D). Notably, the inclusion of an implicit delay in virus production produced a smoother, broader viral load peak, eliminating the unrealistically sharp peak predicted by Model 1 (Fig. 3B).

### Cytokine feedback is necessary to include in the modified target-infected cell model to explain strain-specific viral dynamics

We next investigated whether incorporating positive feedback in IFN production into Model 2 could improve its ability to reproduce the experimental data for the IBV-DMV/1639 strain, as observed with Model 1. Thus, we extended Model 2 by introducing a positive feedback term for cytokine production (Fig. 4A), following the same approach used in Model 1. In particular, we introduced the following dynamics

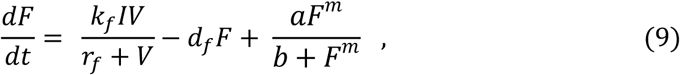

**Figure 4:**
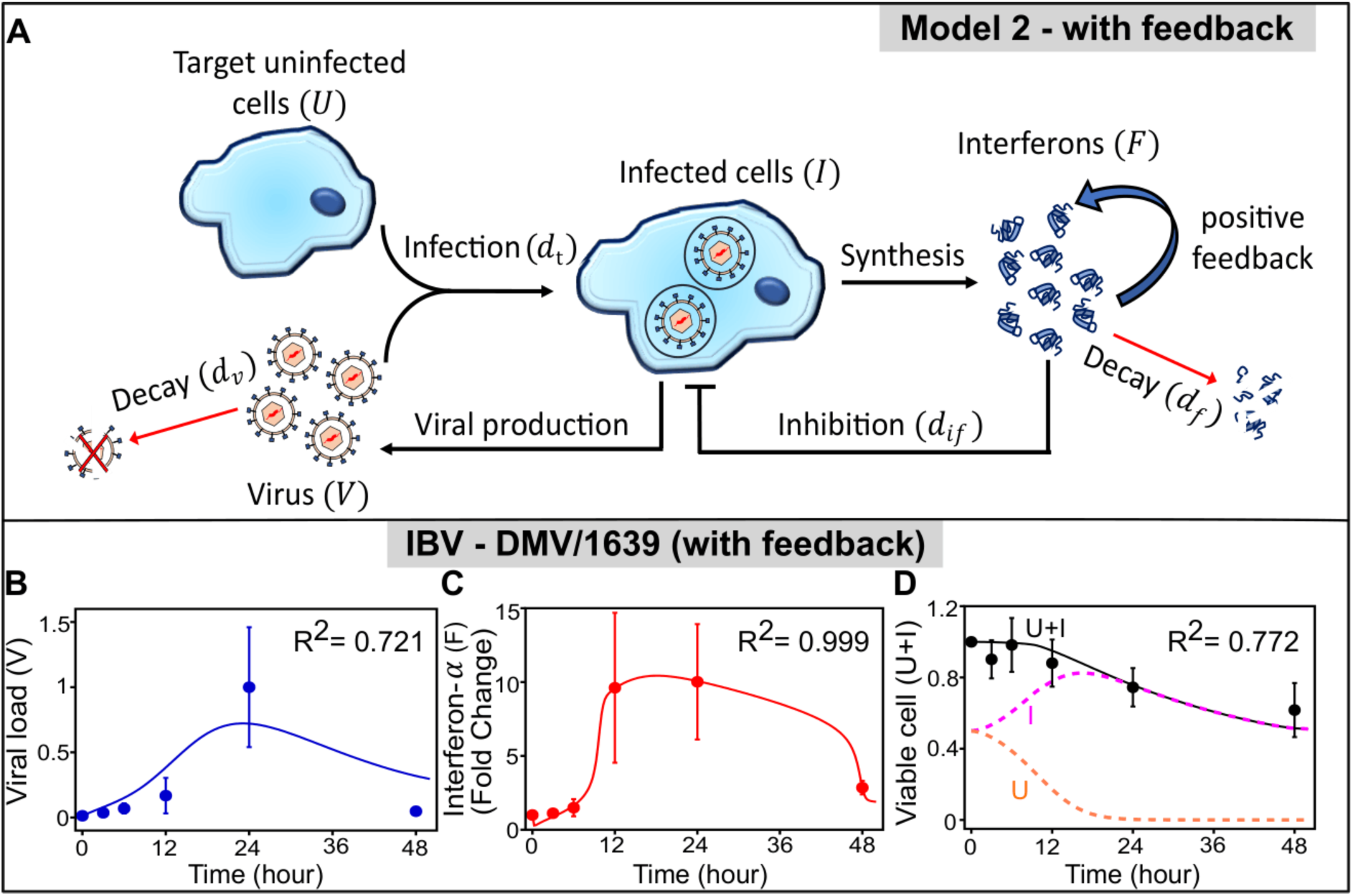
Modified Model 2 with a positive feedback on interferon production captures the infection dynamics of the IBV-DMV strain. **(A)** The schematic of the modified model 2, incorporating an additional positive feedback on the interferon production. **(B–D)** Comparison between model prediction and experimental data for the IBV-DMV/1639 strain with the positive feedback. Panel B presents the viral load trajectory, panel C shows interferon-α concentration levels, and panel D displays the number of viable host cells. Solid black lines correspond to model predictions and filled circles represent experimental data (n = 3; error bars indicate standard deviations). The magenta and orange dashed lines in panel D represent model predictions of infected and uninfected cell populations, respectively.

where, *a* and *b* are the feedback parameters, and for this modified Model 2, we kept the Hill coefficient *m* fixed at 2.

The modified Model 2, described by Eqs. 5, 6, 8, and 9, was solved numerically and fitted to the experimental time-course data for the IBV-DMV/1639 strain (Fig. 4B–4D). The best-fit parameter estimates for both strains, obtained using Model 2, are shown in Table 2. As with Model 1, incorporating positive feedback substantially improved the agreement between model predictions and experimental observations, particularly for the viral-load dynamics (Fig. 4B). These results suggest that cytokine-mediated positive feedback is an important determinant of the infection kinetics of the IBV-DMV/1639 strain. Notably, this conclusion is consistent across both Model 1 and Model 2, indicating that the inferred role of cytokine feedback is robust to the choice of models.

**Table 2.**
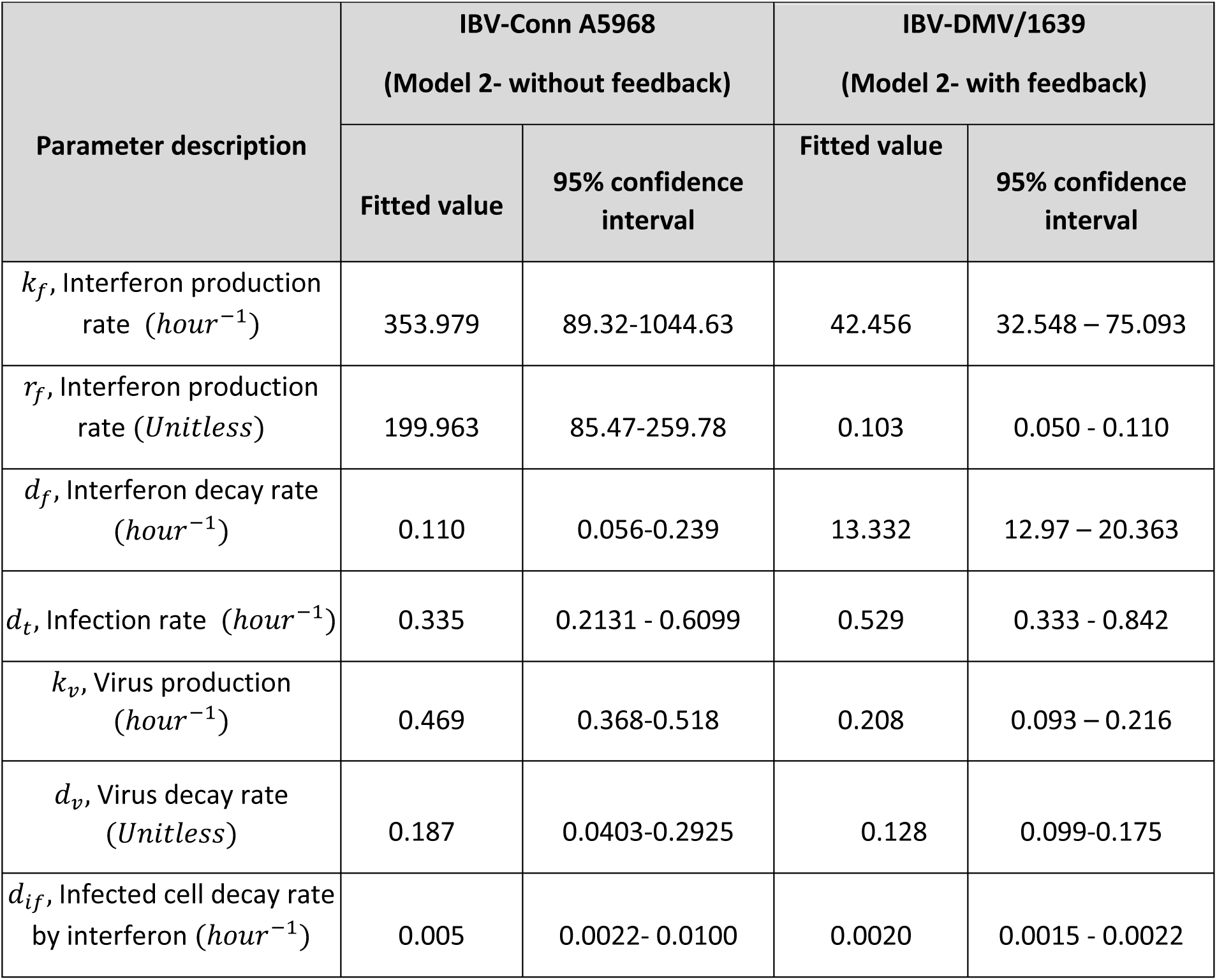
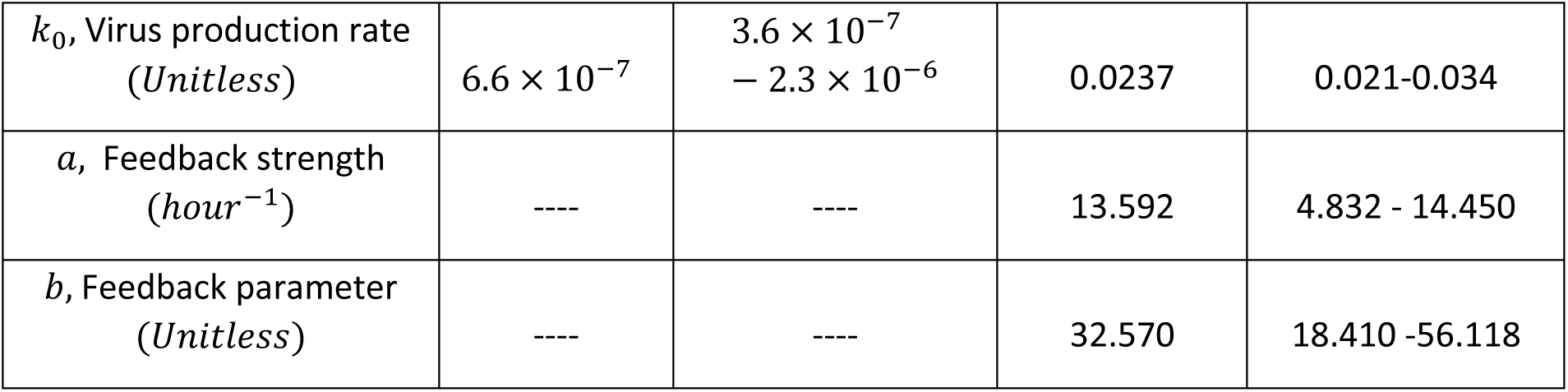
Parameter values estimated from non-linear least squares fitting for IBV-Conn A5968 and IBV-DMV/1639 using Model 2.

| **Parameter description** | **IBV-Conn A5968**<br>**(Model 2- without feedback)** |  | **IBV-DMV/1639**<br>**(Model 2- with feedback)** |  |
|---|---|---|---|---|
|  | **Fitted value** | **95% confidence interval** | **Fitted value** | **95% confidence interval** |
| *k<sub>f</sub>*, Interferon production rate (*hour<sup>−1</sup>*) | 353.979 | 89.32-1044.63 | 42.456 | 32.548 – 75.093 |
| *r<sub>f</sub>*, Interferon production rate (*Unitless*) | 199.963 | 85.47-259.78 | 0.103 | 0.050 - 0.110 |
| *d<sub>f</sub>*, Interferon decay rate (*hour<sup>−1</sup>*) | 0.110 | 0.056-0.239 | 13.332 | 12.97 – 20.363 |
| *d<sub>t</sub>*, Infection rate (*hour<sup>−1</sup>*) | 0.335 | 0.2131 - 0.6099 | 0.529 | 0.333 - 0.842 |
| *k<sub>v</sub>*, Virus production (*hour<sup>−1</sup>*) | 0.469 | 0.368-0.518 | 0.208 | 0.093 – 0.216 |
| *d<sub>v</sub>*, Virus decay rate (*Unitless*) | 0.187 | 0.0403-0.2925 | 0.128 | 0.099-0.175 |
| *d<sub>if</sub>*, Infected cell decay rate by interferon (*hour<sup>−1</sup>*) | 0.005 | 0.0022- 0.0100 | 0.0020 | 0.0015 - 0.0022 |
| *k<sub>0</sub>*, Virus production rate (*Unitless*) | 6.6 × 10<sup>−7</sup> | 3.6 × 10<sup>−7</sup> − 2.3 × 10<sup>−6</sup> | 0.0237 | 0.021-0.034 |
| *a*, Feedback strength (*hour<sup>−1</sup>*) | ---- | ---- | 13.592 | 4.832 - 14.450 |
| *b*, Feedback parameter (*Unitless*) | ---- | ---- | 32.570 | 18.410 -56.118 |

### Data-driven inference and validation of the interferon feedback

We next used a data-driven, physics-informed machine-learning (ML) framework (42, 43) to infer the presence and strength of feedback in cytokine dynamics directly from the experimental data. Using Model 1 as the underlying mechanistic framework, we generated approximately 10,000 bootstrap samples at each experimentally measured time point (3, 6, 12, 24, and 48 h). All functional terms and parameters in Model 1 were treated as known, except for the feedback term in Eq. 4. We then computed weak-form integral residuals from the bootstrap datasets, using the corresponding state variables (see Materials and Methods). These residuals were used to identify the unknown feedback contribution, which was assumed to follow a Hill-type functional form. By constructing a library of Hill functions with varying parameters and applying linear least-squares regression, we inferred the feedback parameters *a*, *b*, and *m*. The bootstrap datasets were then partitioned into training and test sets using a 70:30 split, and the feedback parameters were estimated from the training data for both IBV-Conn A5968 and IBV-DMV/1639 (Fig. 5A and Table S1). The inferred feedback was negligible for IBV-Conn A5968, whereas IBV-DMV/1639 consistently exhibited strong Hill-type positive feedback, with an inferred Hill exponent of *m* = 2.

**Figure 5.**
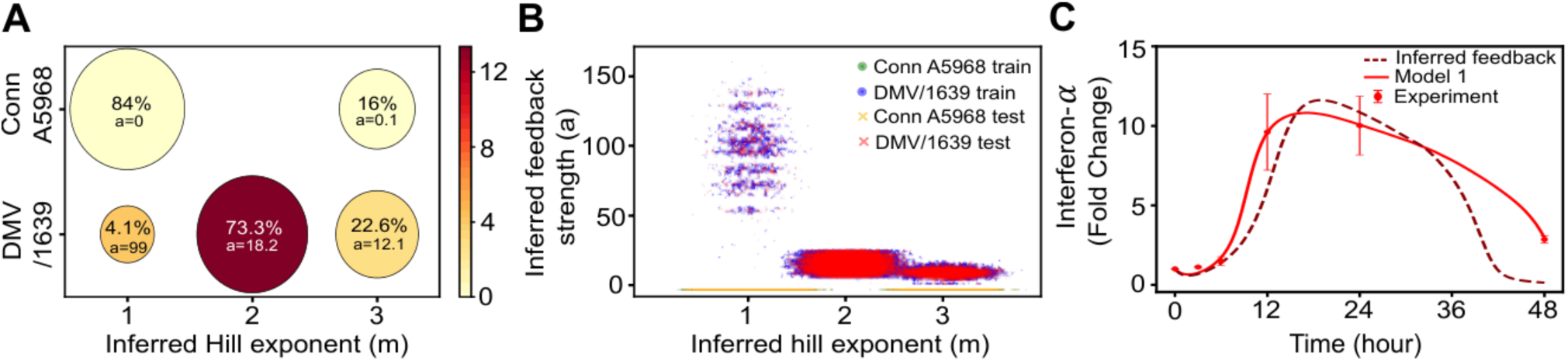
Identification and validation of inferred interferon feedback dynamics using a physics-informed ML framework. **(A)** Summary of the inferred interferon feedback parameters obtained from the training datasets for IBV-Conn A5968 and IBV-DMV/1639 strains. The color bar represents the weighted feedback strength, i.e., the fraction of datasets with the corresponding inferred Hill exponent *m* out of all the training datasets times the mean feedback strength, *a*. For IBV-Conn A5968, regardless of the Hill exponent, the inferred feedback strength is approximately zero, indicating negligible feedback. For most of the IBV-DMV/1639 training datasets (73.3%), the inferred Hill exponent is 2, with a mean inferred feedback strength of approximately 16.8, supporting the existence of a Hill-type feedback. **(B)** Classification using k-nearest neighbors (kNN) algorithm, showing the distribution of training and test samples based on the inferred Hill exponent and feedback strength. The scatterplot indicates that the two viral strains are well separated in their feature-space with little-to-no overlap. **(C)** Comparison of Model 1 with feedback and the inferred feedback model against the experimental data of interferons for the IBV-DMV/1639 strain, illustrating the agreement of ML-based inference with both the mathematical model and the data.

We next applied a k-nearest neighbors (kNN) classifier (44) to the test datasets and visualized the results in the feature space defined by the inferred Hill exponent, *m* and feedback strength, *a* (Fig. 5B). These inferred parameters thus served a dual purpose: quantifying the feedback dynamics and providing features for distinguishing between the two viral strains. The resulting feature-space distributions showed clear separation between IBV-Conn A5968 and IBV-DMV/1639, with almost no overlap.

Finally, we incorporated the ML-inferred feedback parameters into Model 1 and compared its predictions with those of the reference model fitted to the experimental IFN time-series data for IBV-DMV/1639 (Fig. 5C). Because the inferred feedback was negligible for IBV-Conn A5968, the ML-inferred and reference models were effectively identical for this strain. For IBV-DMV/1639, the ML-inferred model reproduced the observed IFN dynamics with reasonable agreement (normalized root-mean-square error, NRMSE = 0.195 for the ML-inferred model versus 0.020 for the reference Model 1). Thus, the ML framework independently identified strain-specific feedback dynamics and recovered the principal features of the observed IFN response.

## Discussion

Of late, mathematical models of varying complexity have become valuable tools for understanding viral infection dynamics within hosts (11, 12, 45, 46). However, uncertainties in the underlying biology, particularly the interactions among viruses, host cells, and immune mediators, as well as limited experimental data, often constrain model validation and parameter estimation. These challenges are particularly relevant for avian viral pathogens, for which quantitative datasets capturing both infection kinetics and host immune responses remain relatively limited.

In this study, we revisited and extended classical within-host viral dynamic models to characterize the infection kinetics of IBV in chicken macrophages. Despite their relative simplicity, these models captured key features of viral growth and decline and provided mechanistic insights into the processes shaping infection dynamics. We found that the IBV-Conn A5968 strain could be adequately described by a relatively simple model framework (Model 1 in Fig. 1), whereas the IBV-DMV/1639 strain required the inclusion of a Hill-type positive feedback term in IFN production (Fig. 2). The Hill exponent (*m*), which controls the degree of nonlinearity in the feedback response, was an important determinant of the DMV/1639 dynamics. A Hill exponent of *m* = 1 failed to adequately reproduce the experimental data (Fig. S2), whereas *m* = 2 substantially improved the fit (Fig. 2). In contrast, the fit for IBV-Conn A5968 was largely insensitive to the choice of *m*, consistent with the negligible contribution of positive cytokine feedback in this strain (Fig. S2). Importantly, our data-driven, physics-informed machine-learning framework independently inferred *m* = 2 for IBV-DMV/1639 strain (Fig. 5A), providing an independent validation of the nonlinear feedback identified by the mechanistic model. Together, these results suggest that the two IBV strains differ in the regulatory mechanisms governing their interactions with the host antiviral response.

We further evaluated the robustness of the inferred mechanisms using two modelling frameworks of increasing complexity (Models 1 and 2) and reported the best-fit parameter values from these models (Tables 1 and 2). To quantify uncertainty in parameter estimation, we performed bootstrap resampling by repeatedly generating resampled datasets from the original experimental data and refitted the models to each resampled dataset. The resulting parameter distributions were generally well constrained, indicating that the inferred biological mechanisms were not dependent on a particular realization of the experimental data (Figs. S6–S9). The corresponding 95% confidence intervals provide a robust quantitative estimates of the uncertainty associated with each parameter (Tables 1 and 2).

A central finding of this study is the inferred positive feedback in IFN production for the IBV-DMV/1639 strain. The biological significance of this feedback can be appreciated from the early IFN dynamics (compare Fig. 1C and 1F). Following infection, IFN-α levels increased more rapidly for IBV-DMV/1639, reaching approximately 10-fold above the initial level by 12 h post-infection. In contrast, IFN-α increased more gradually for IBV-Conn A5968, reaching a similar magnitude only by 24 h. This difference in the temporal dynamics is consistent with the stronger nonlinear feedback inferred for IBV-DMV/1639 by both the mathematical modelling and ML approaches. However, our study identifies the feedback phenomenologically and does not establish the underlying molecular pathway. One plausible mechanism involves the regulatory network controlling type I IFN production. In mammalian systems, viral infection activates interferon regulatory factor 3 (IRF3), which contributes to type I IFN production and activation of the JAK–STAT pathway (30). This pathway induces interferon-stimulated genes (ISGs), including IRF7, which can further enhance transcription of type I IFN genes and thereby establish a positive-feedback loop (23). However, this canonical IRF3–IRF7 mechanism cannot be directly extrapolated to chickens, where a true IRF3 ortholog has not been identified (47). Functional studies instead indicate that IRF7 is a major regulator of type I IFN production in chickens and can mediate IFN-β responses, suggesting that IRF7 may partially compensate for the absence of IRF3 (48). Also, overexpression of chicken IRF7 has been shown to enhance IFN-α and IFN-β expression following viral stimulation [4]. Furthermore, a recent study reported that IBV infection can indirectly enhance IFN-β production in chicken cells following stimulation of MDA5- or TLR3-dependent pathways, producing a positive-feedback-like amplification of the IFN response (49).

Our work therefore provides a quantitative starting point for investigating the molecular basis of the identified IFN feedback. Future experiments targeting candidate regulators such as IRF7, MDA5, TLR3, and downstream signaling components could test whether these pathways account for the inferred feedback and its differences between IBV strains. Such experimental work could ultimately connect the phenomenological feedback term in our present models to specific molecular mechanisms. Moreover, extending our model to a stochastic framework (50, 51) can be a useful future direction, since large cell-to-cell variability can be observed in some viral infections (52).

## Acknowledgments

KM acknowledges the University Grants Commission (UGC), Government of India, for the PhD fellowship. DD acknowledges the SERB-SIRE Fellowship (SIR/2022/000156) for supporting a collaborative visit to the University of Calgary, where this work was conceived, and financial support from SERB/ANRF (EEQ/2023/000551 and ANRF/ARGM/2025/001984/TS). DD also thanks Muhammad Farooq (University of Calgary) and Prof. Amirul I. Mallick (IISER Kolkata) for initial discussions.

## Supporting information

Supporting Information contains Materials and Methods, 9 supplemental figures, and supplemental tables.

## Materials and methods

### Experimental Methods

#### Infection of avian macrophages with IBV

As previously described (1, 2), Macrophages (MQ-NCSU cells) were suspended in LM-Hahn medium supplemented with 8% fetal bovine serum (FBS), 10% chicken serum, 1% antibiotic–antimycotic solution, and 10 µM 2-mercaptoethanol, and seeded into 6-well plates at a density of 1.5×10⁶ cells per well, all cell culture reagents were purchased from Gibco Life Technologies (Burlington, ON, Canada). The cells were then incubated at 40°C in 5% CO₂ for 24 hours. Once a confluent monolayer was established, the cells were washed twice with Dulbecco’s phosphate-buffered saline (dPBS) and infected in triplicate with IBV DMV/1639 or Conn A5968 strains at a multiplicity of infection (MOI) of 0.1, or mock-infected as controls. After a one-hour adsorption period, the inoculum was completely removed, and the cells were washed twice with dPBS (Sigma-Aldrich, Burlington, MA, USA ) to eliminate any residual virus. The cells were then incubated with the same serum-free LM-Hahn medium (Gibco Life Technologies, Burlington, ON, Canada) at 37°C in 5% CO₂. At 3, 6, 12, 24, and 48 hours post-infection (hpi), RNA extracted from cell culture supernatants was used to quantify IBV RNA levels. The experiment was performed in triplicate and repeated three times for reproducibility.

#### Type 1 interferon (IFN) activity during macrophage infection

To determine whether type 1 IFN activity is induced in macrophages following IBV infection in vitro, MQ-NCSU cells were seeded in 12-well plates at a density of 0.75×10⁶ cells per well and incubated for 24 hours. The cells were then either infected with 0.1 MOI of IBV DMV/1639 strain or 0.1 MOI of IBV Conn A5968 strain, or treated with 50 μg/ml double-stranded (ds) RNA (Invivogen, San Diego, California, USA) as a positive control for type 1 IFN induction. RPMI medium was used as the negative control. All experiments were performed in triplicate and repeated three times. At the previously indicated time points, culture supernatants were collected and stored at -80°C for subsequent quantification of type 1 IFN activity.

#### RNA extraction and complementary DNA (cDNA) synthesis

Trizol LS reagent (Ambion, Invitrogen Canada Inc., Burlington, ON, Canada) was used to extract RNA from 250 μl of macrophage-culture supernatants, following the manufacturer’s instructions. The RNA pellet was resuspended in 20 μl of RNase-free water, and its concentration was measured using a Nanodrop 1000 spectrophotometer at 260 nm (Thermo Scientific, Wilmington, DE, USA). A total of 2000 ng of RNA was transcribed into cDNA using 10X RT random primers and the High Capacity cDNA Reverse Transcription Kit (Invitrogen Life Technologies, Carlsbad, CA), according to the manufacturer’s instructions.

#### IBV RNA quantification

A quantitative PCR (qPCR) assay was performed to quantify IBV RNA in cell culture supernatants. The assay used gene-specific primers targeting the IBV N gene: forward primer IBV-N Fw (5’-GACGGAGGACCTGATGGTAA-3’) and reverse primer IBV-N Re (5’-CCCTTCTTCTGCTGATCCTG-3’) (3). All qPCR reactions were conducted using a CFX96 Real-Time System C1000 Thermal Cycler (Bio-Rad Laboratories, Mississauga, ON) in 96-well un-skirted, low-profile PCR plates (VWR, Edmonton, AB, Canada), sealed with MicroAmp Optical Adhesive Films (Applied Biosystems, Foster City, CA, USA).

Each reaction was performed in a final volume of 20 μL, consisting of Fast SYBR™ Green Master Mix (Invitrogen, Burlington, ON, Canada), 5 nM of each primer (3), 200 ng of cDNA derived from each sample, and RNAse-free water. To ensure accuracy, the qPCR assay included a positive control (IBV N-gene plasmid), a negative template control (NTC) using RNAse-free water, and a no reverse transcriptase control (NRT) to rule out genomic DNA contamination. A standard curve was generated using a serial dilution of the IBV N-gene plasmid, as previously described (3). The oligonucleotide primers were synthesized by Integrated DNA Technologies (IDT, Coralville, IA, USA)

The qPCR thermal cycling conditions were as follows: an initial denaturation step at 95°C for 20 seconds, followed by 40 amplification cycles of 95°C for 3 seconds and 60°C for 30 seconds. A melting curve analysis was performed with the following steps: 95°C for 10 seconds (Segment 1), 65°C for 5 seconds (Segment 2), and 90°C for 5 seconds (Segment 3), with fluorescence acquisition at 60°C for 30 seconds.

To determine amplification efficiency, a standard curve was generated across a linear dynamic range (LDR) spanning 7 log₁₀ to 1 log₁₀ copies per reaction. The efficiency and linearity of the qPCR assay were verified by assessing the slope and correlation coefficient (R²) of the standard curve, ensuring reliable and reproducible quantification of IBV RNA.

#### Estimation of percentage of dead MQ-NCSU cells in culture

After collecting the MQ-NCSU culture supernatants, the adherent cells were scraped into fresh media and resuspended. The cell suspensions were then centrifuged at 1000 rpm for 10 minutes at 4°C. The resulting cell pellet was resuspended in 100 μL of growth media and kept on ice until cell counting. Cell viability was assessed using a hemacytometer following Trypan blue staining (Gibco Life Technologies, Burlington, ON, Canada), with both live and dead cells enumerated. The percentage of dead cells in each well was calculated relative to the total cell count in the respective well.

#### Extracellular IFN-α ELISA

Chicken interferon-alpha (IFN-α) levels in MQ-NCSU cell culture supernatants were measured using a Chicken IFN-alpha ELISA Kit (Abcam, Cambridge, UK; catalog no. ab273260), according to the manufacturer’s instructions. The assay included blank wells, standards, and positive and negative controls. Absorbance was measured at 450 nm using a microplate reader (680 XR, Bio-Rad Laboratories, Hercules, CA, USA), and IFN-α concentrations in the culture supernatants were calculated from the generated standard curve.

## Numerical Methods

### Normalization of experimental data for comparing with Model predictions

#### Viable Cells

The number of viable cells is normalized by the initial cell count at time zero, and was expressed as the fraction of the initial cell number. In experiments, the initial cell counts were kept at 1.5 × 10^6^. Accordingly, in mathematical models, the total cell numbers (T) were taken as a normalized variable such that the initial fraction of viable cells is T(0)=1.

#### Extracellular Virus

The extracellular virus concentration is normalized by the maximum value of the mean, *V^max^* (i.e., the peak value), measured across all time points. This ensures that both experimental and simulated viral dynamics are represented on a comparable relative scale (see the initialization for mathematical models below).

In general, viral RNA copy numbers measured by RT-qPCR may overestimate the concentration of infectious viral particles, typically quantified as plaque-forming units (PFU/mL), because RT-qPCR can detect viral RNA from both infectious and non-infectious sources. Nevertheless, we use the viral RNA copy number, normalized by the maximum mean copy number observed across the dataset, as a proxy for viral load.

#### Extracellular IFN-α

In mock cultures (i.e., cultures without viral exposure), experimental data showed that IFN-α protein levels remained relatively constant, indicating a basal expression level. Thus, the IFN-α values in experiments are normalized to the IFN level observed in the mock condition. In models, we assume that the initial IFN-α expression is the basal level, and hence we normalize by this variable by the initial value. Accordingly, in the models, the IFN-α level (F) was expressed as a normalized variable, with the initial IFN-α level F(0)=1.

### Initial conditions for the mathematical models

#### Initial extracellular viral load

The initial cell count in the experiments was 1.5 × 10^6^ with a multiplicity of infection (MOI) of 0.1, yielding the total initial viral load as: *V^experimental^*(0) = MOI × initial cell number = 1.5 × 10^5^. We assume that a fraction of this initial viral population leads to infection, while the rest remains in the extracellular space at time zero. As described in the data normalization section, extracellular viral load is normalized by the peak of mean viral load of each strain, *V^max^*. Thus, the initial value of the normalized extracellular virus in the both Model 1 and 2 were taken as

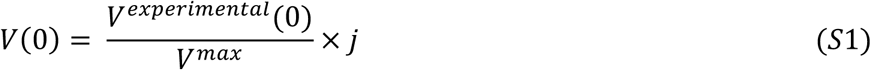

where the fraction *j* was taken as 0.5 in the main text, and was also varied (*j* = 0.6, 0.7, 0.8) in the SI (Figs. S1, S3, S4, and S5).

#### Initial viable cell

In Model 1, the target cell population is set to 1 at time zero (i.e., *T*(0) = 1) since *T*(*t*) was taken as a normalized variable in our model. However, for Model 2, total target cell populations were categorized into two populations: uninfected and infected, i.e., *T*(*t*) = *U*(*t*) + *I*(*t*). We assume in Model 2 that the relative abundance of uninfected and infected cells follows the same ratio as that of extracellular to intracellular virus. Thus, the initial fractions of uninfected and infected cells were assumed to be *U*(0) = *j* and *I*(0) = 1 − *j*, respectively, such that the total cell fraction remains conserved, i.e, *T*(0) = *U*(0) + *I*(0) = 1.

#### Initial IFN-α level

Since the IFN-α level is expressed as a normalized variable relative to its initial value, the IFN levels are set to 1 at time zero for both Model 1 and 2 (i.e, *F*(0) = 1).

#### Best-fit parameter values and data fitting using least squares optimization method

We fixed some parameters and estimated the rest by fitting the model to the experimental data of viral load, IFN fold change, and percentage of viable cells. The feedback hill coefficient (*m*) in IFN equation (Equations 2 and 7) was fixed at 2 for both models. The initial population of the cell and viral load were set at 1.5 × 10^6^ and 1.5 × 10^5^ ,respectively. The initial IFN fold change was set to 1, indicating no change at the starting point. In the three-variable model, all cells were assumed to be target cells capable of producing virus. In the four-variable model, the initial populations of infected and uninfected cells were assumed to be equal, such that their sum equal to the total initial cell population. The remaining parameters were estimated from data fitting. We fit the models to the viral load, IFN data and cell data of two different strain (DMV/1639 and Conn A5968) of IBV virus using the python package ‘curve_fit’ (version 3.8). Parameter estimation was performed by fitting the model equations to the experimental data using a nonlinear least squares optimization method. The program minimized the sum of squared residual(SSR) between data point and corresponding model prediction, given by

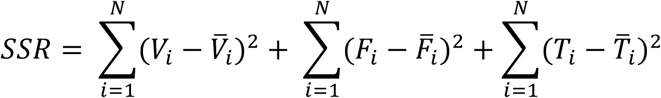

Here, *N* denotes the total number of data points for each variable, which is identical across all variables. The experimental viral load data are denoted by *V_i_* , with the analogous model predictions represented by *V*^;^*_i_* . Similarly, the measured IFN fold change and the number of living cell are denoted by *F_i_* and *T_i_* respectively, while the corresponding model outputs are represented by *F*^;^*_i_* and *T*^;^*_i_* respectively. In the four variable model, *T*^;^*_i_* is the sum of uninfected cell and infected cell. The optimal parameter set was identified as the one that provided the maximum coefficient of determination (*R*^2^).

#### Parameter distribution and confidence interval

To evaluate uncertainty of model parameters, we applied a bootstrap resampling strategy using the MATLAB ‘bootstrp’ package. A total of 100,000 bootstrap datasets were generated from the original experimental data. For each resampled dataset, model parameters were estimated using the nonlinear least-squares fitting function ‘curve_fit’ from the SciPy library in Python. To ensure the reliability of parameter estimates, only those parameter sets that achieved a *R*^2^ value greater than 0.8 were retained for further analysis.

The distributions of the selected parameters were subsequently characterized by fitting them to either gamma or exponential distributions using MATLAB’s built-in ‘fit’ functions. From these fitted distributions, 95% confidence intervals for each parameter were computed. This approach enabled us to quantify parameter uncertainty with respect to experimental variability.

### Supervised Machine Learning framework to infer IFN feedback

#### Construction of weak-form integral residuals

We consider the interferon (IFN-α) equation of Model 1 (Eq. 4, main text) and assume that all terms except the positive feedback term *P*(*F*) are known. The bootstrap trajectories provide the state variables, while previously estimated best-fit model parameters are used for all known coefficients (Table 1, main text). Rearranging the equation to have all the known terms on one side, we get:

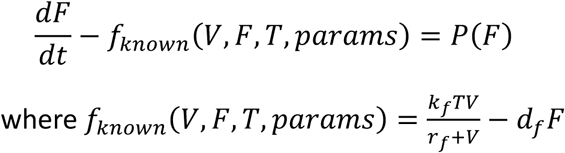

For consecutive experimental time points *t_k_* and *t_k_*_+1_, we integrate the above equation:

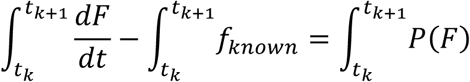

Using the trapezoidal rule, we can approximate this as:

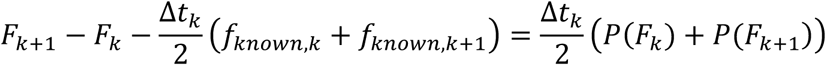

The left-hand side is defined as the weak-form integral residual (4):

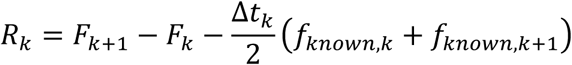

which captures the component of the interferon (IFN-α) dynamics not explained by the known model terms. If positive feedback exists, it must account for the structure of *R_k_*. For each trajectory, the residuals over all observation intervals are collected into a residual vector **R**.

#### Hill function library construction for the feedback term

We assume the feedback term to take the form of a Hill function:

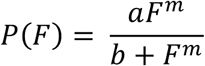

Rather than fixing *m* = 2 *a priori*, we considered *m* ɛ {1,2,3}. Depending on the value of *m*, we fixed a range of *b* values as follows:

For *m* = 1 : 0.1 ∗ *F*_min_ ≤ *b* ≤ 10 ∗ *F*_max_ as a biologically feasible range

For 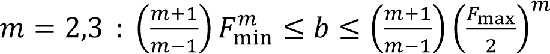 such that the inflection point of the Hill function lies in the range [*F*_min_, *F*_max_/2] ensuring that the transition region of the Hill function lies within the observed range of IFN-α concentrations. Since parameter inference is performed independently for each bootstrap trajectory, the admissible range of *b*is computed separately for each trajectory using its corresponding *F*_min_ and *F*_max_.

Using the above, a grid of (*m*, *b*) values is generated. Each (*m*, *b*) pair corresponds to a unique Hill function.

#### Regression-based identification

For a fixed (*m*, *b*) pair, we numerically integrate the Hill function according to the trapezoidal rule to get:

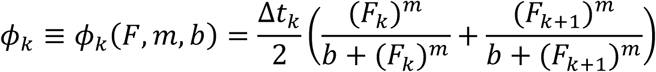

These are also organized into a vector **φ** for all the trajectories and time-points. The problem now becomes linear in *a*:

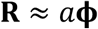

For every candidate pair (*m*, *b*), the optimal value of *a* is obtained by linear least-squares regression. The parameter combination producing the smallest mean squared error is selected as the inferred feedback model. A non-zero inferred value of *a* indicates evidence for positive IFN-α feedback.

#### Model training: extracting features for each viral strain

The time-series data for both the viral strains are split into training and test sets in a 70:30 ratio. The above protocol is implemented on the training datasets to extract the feature vector *θ* = (*a*, *b*, *m*) for each individual time-series for each strain. The inferred parameter vector *θ* from each training trajectory was used as the feature representation for subsequent classification.

#### Test set classification

Next, the test set trajectories for both the strains are combined and shuffled. The same regression procedure is applied to every individual test trajectory, inferring a θ*_test_* for each trajectory.

Before classification, the features are scaled, i.e., each feature dimension (log(*a*) , log(*b*) , *m*) is re-centered to mean=0 and unit variance. This ensures that no single feature dominates due to scale. Classification is done using *k*-nearest neighbors (5) with *k* = 5.

### ODE model simulation and validation

Finally, we validate the results by simulating the ODE-based Model 1 (Eq. 1, 3, and 4 in the main text) using the inferred parameters and comparing the fit to both the experimental data and the results from the estimated best-fit parameters. The initial values for the state variables *T*, *F*, *V* are taken as the mean values of the experimental data at the first time-point. To quantify and compare the fits, we compute the normalized root-mean-squared error (NRMSE) between the experimental data and the simulations using both the inferred and known parameters.

---

**Figure S1.**
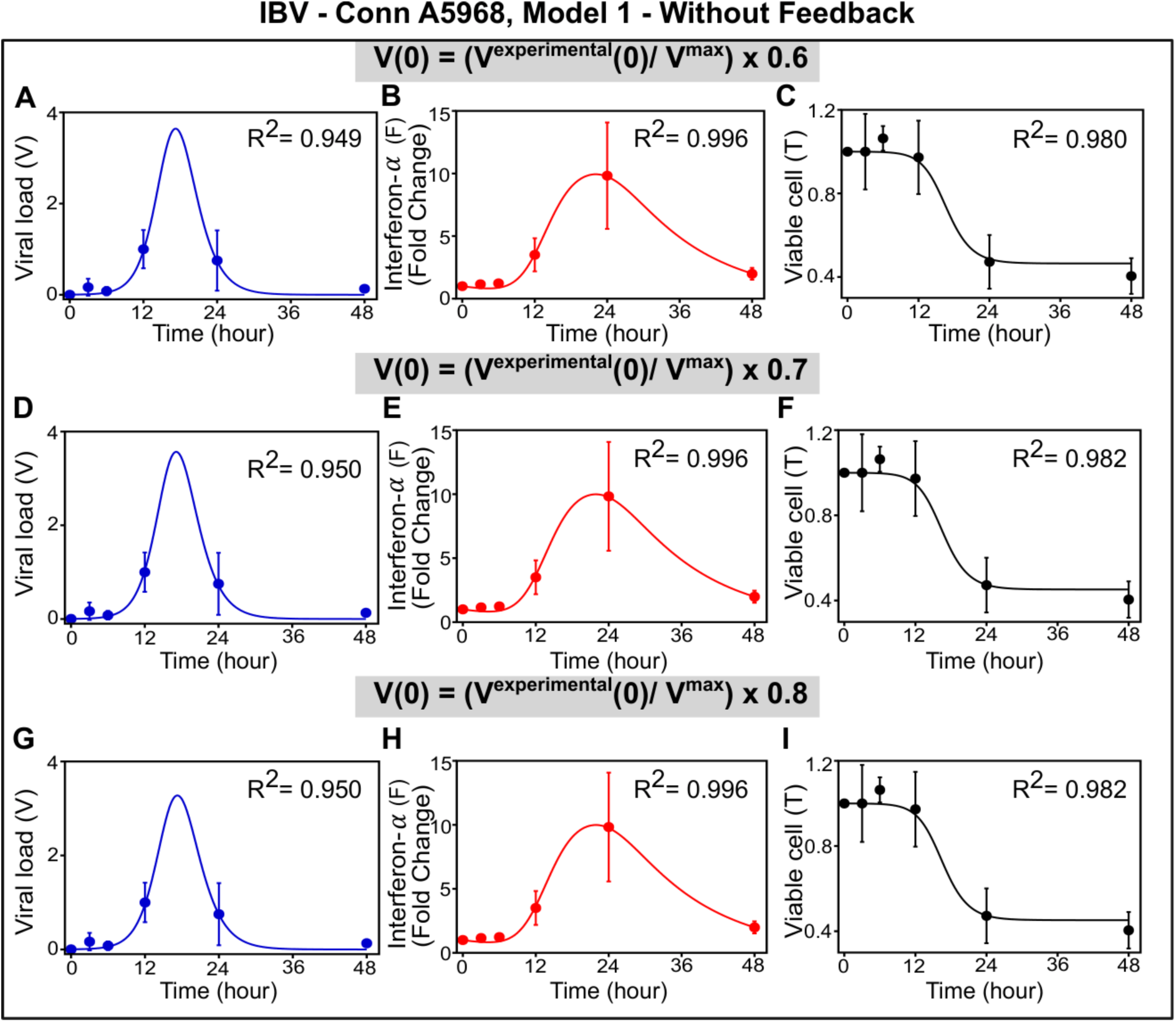
Effect of varying the fraction of initial extracellular virus in Model 1 for the IBV-Conn A5968 strain. Model-predicted temporal dynamics of extracellular viral load, interferon-α, and viable cells are shown by varying the fraction of the initial extracellular viral particles that can infect the cells. The fractions were taken as 0.6 (A-C), 0.7 (D-F), and 0.8 (G-I), while the main figures were shown for a fraction of 0.5 --- see Equation S1 in the SI for the initialization of the Model. Solid lines represent model predictions, and filled circles indicate experimental measurements (n = 3; error bars indicate standard deviation).

**Figure S2.**
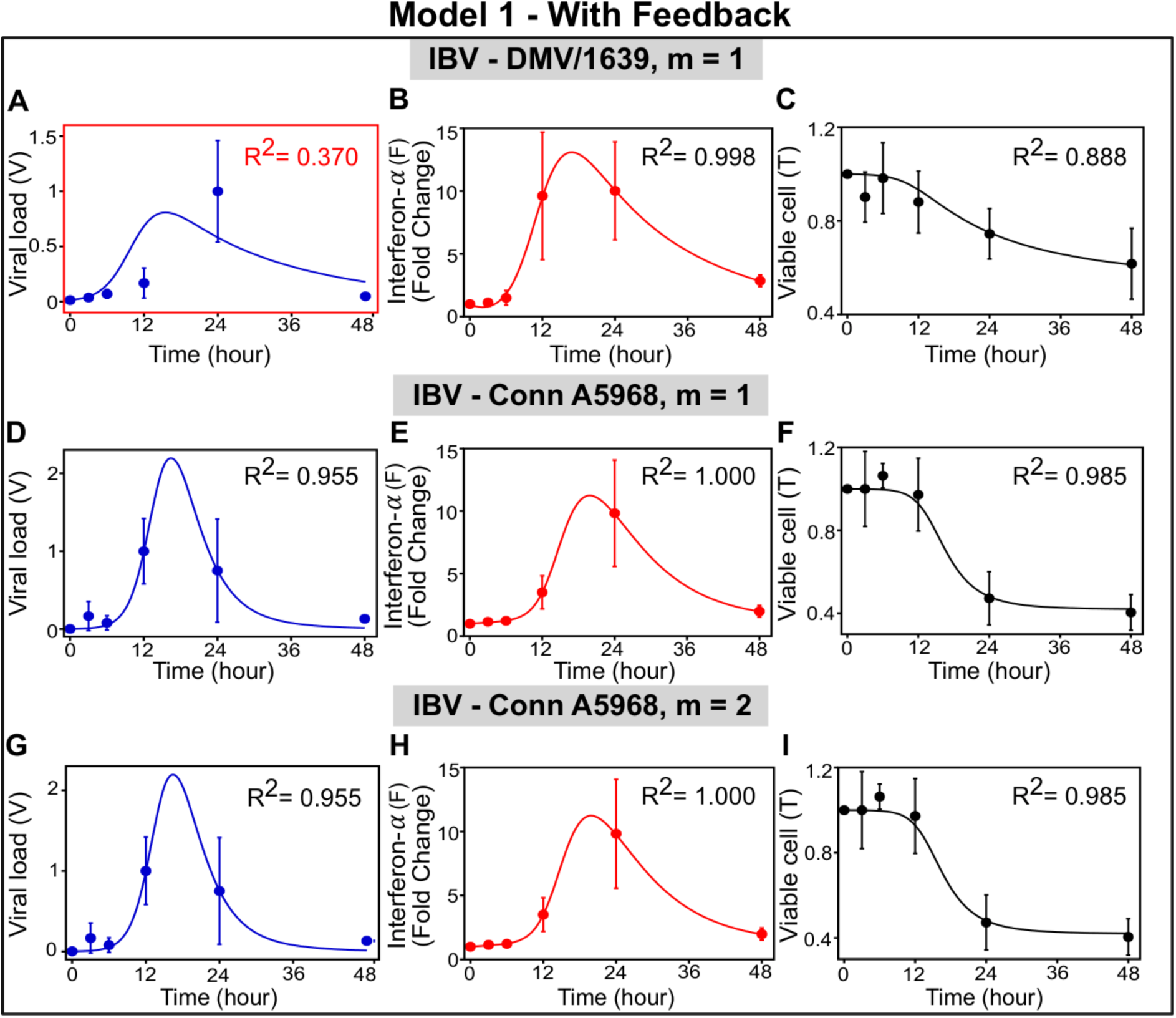
Effect of varying the Hill exponent (m) in Model 1 with the cytokine feedback. Comparison of data and model-predicted temporal dynamics of extracellular viral load, interferon-α, and viable cells for the IBV-DMV/1639 strain with the hill exponent m=1 (A-C), and for the IBV-Conn A5968 strain with m=1 (D-F) and m=2 (G-I). Note that m=1 cannot fit the viral load of the IBV-DMV/1639 strain, while the prediction for m=2 fits well with the data, as given in Fig. 2. However, for IBV-Conn A5968, the model predictions for m=1 and m=2 both match the data but with negligible feedback parameter (b∼10^-9^), suggesting that feedback has no effective role for this strain.

**Figure S3.**
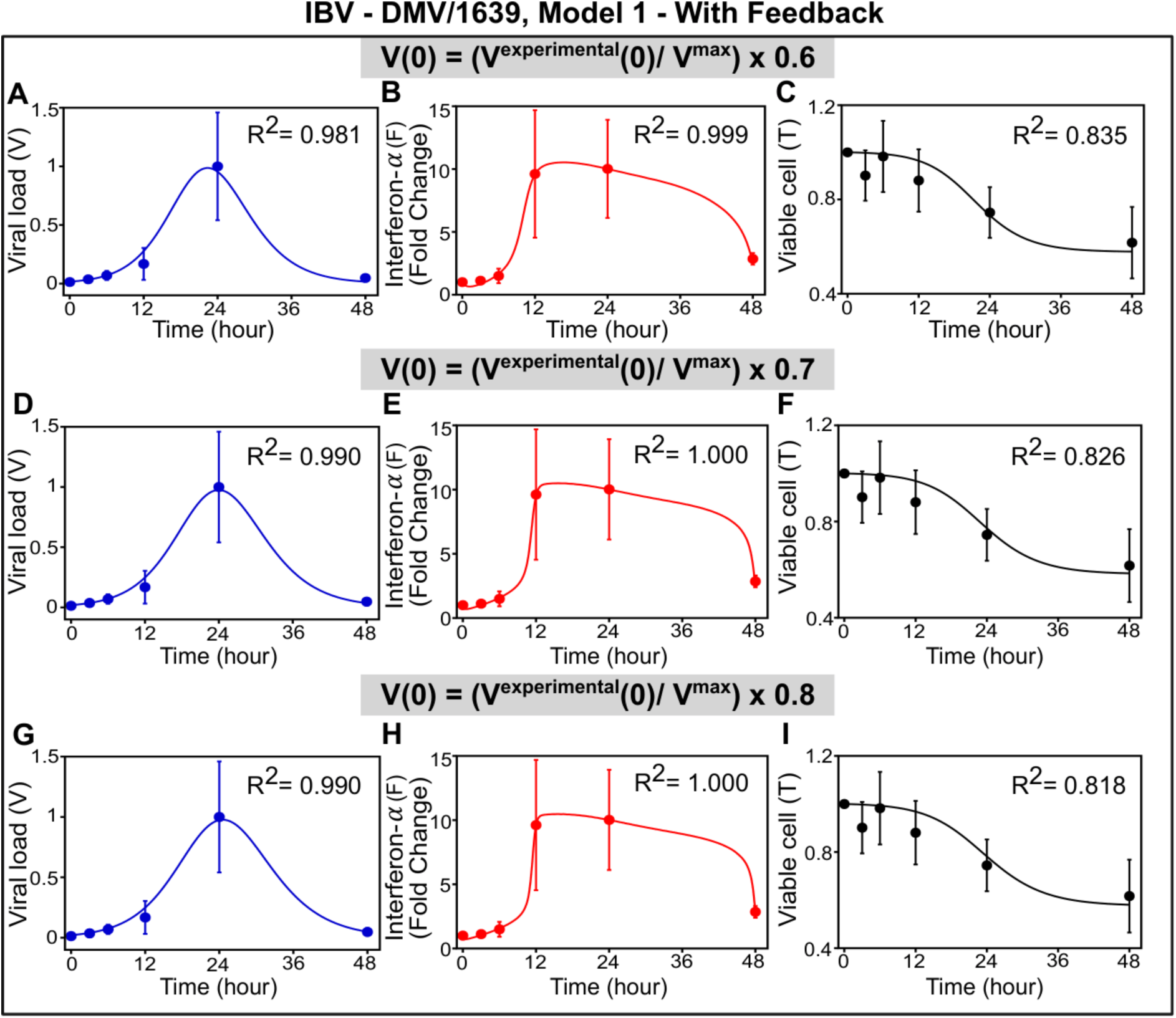
Effect of varying the fraction of initial extracellular virus in Model 1 (with feedback) for the IBV-DMV/1639 strain. Predicted dynamics of extracellular viral load, interferon-α, and viable cells are shown by varying the fraction of the initial extracellular viral particles that can infect the cells. The fractions were taken as 0.6 (A-C), 0.7 (D-F), and 0.8 (G-I), while the main figures were shown for a fraction of 0.5 --- see Equation S1 in the SI for the initialization of the Model. Solid lines represent model predictions, and filled circles indicate experimental measurements (n = 3; error bars indicate standard deviation).

**Figure S4.**
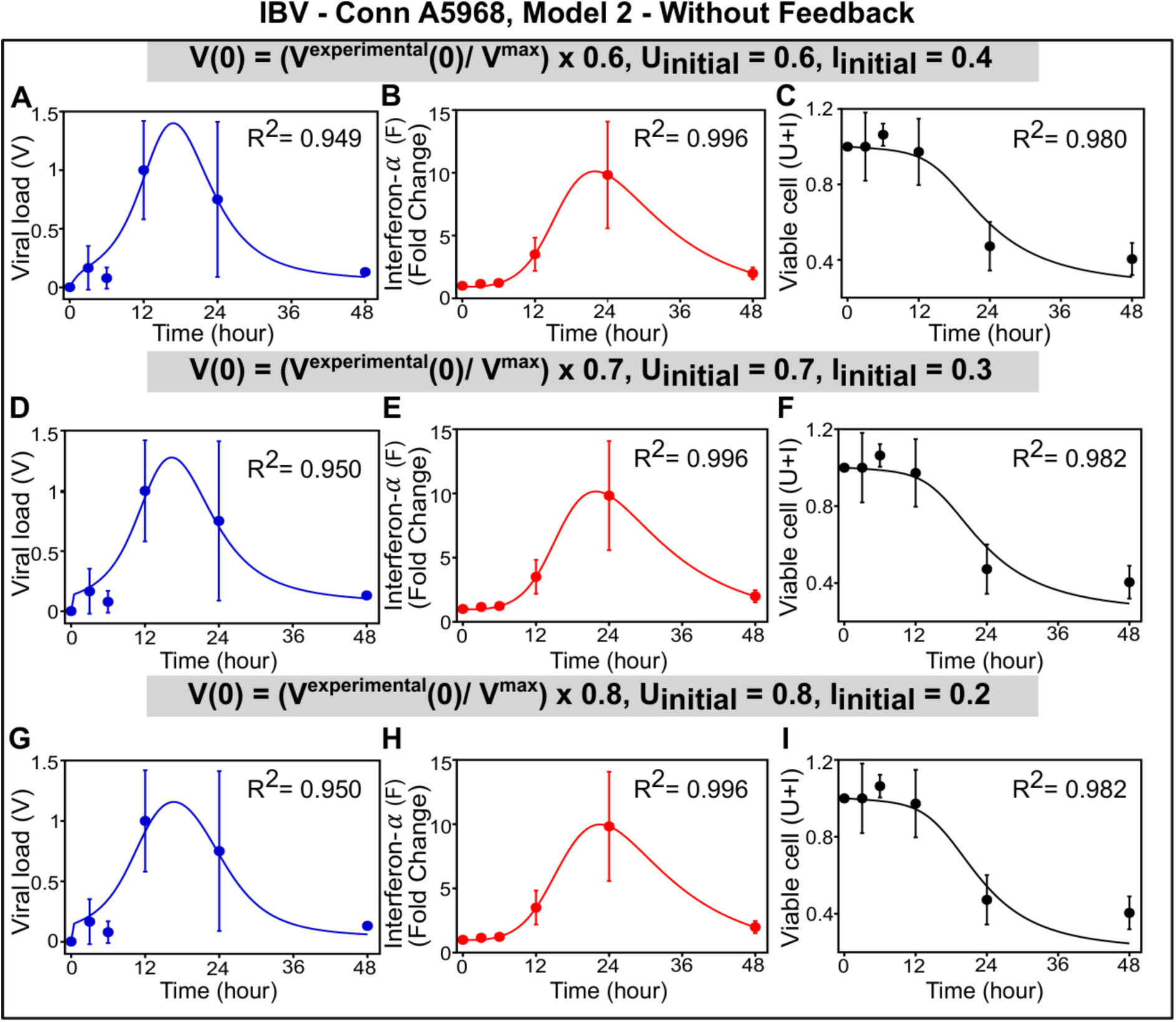
Effect of varying the fraction of initial extracellular virus and the fraction of uninfected and infected cells in Model 2 (for the IBV-Conn A5968 strain). Model-predicted temporal dynamics of extracellular viral load, interferon-α, and viable cells are shown by varying the fraction of the initial extracellular viral particles that can infect the cells. Note that the initial fraction of uninfected cells was taken to be the same as the initial extracellular virus fraction. These fractions were taken as 0.6 (A-C), 0.7 (D-F), and 0.8 (G-I), while the main figures were shown for a fraction of 0.5 --- see Equation S1 in the SI for the initialization of the Model. Solid lines represent model predictions, and filled circles are experimental data (error bars indicate standard deviation).

**Figure S5.**
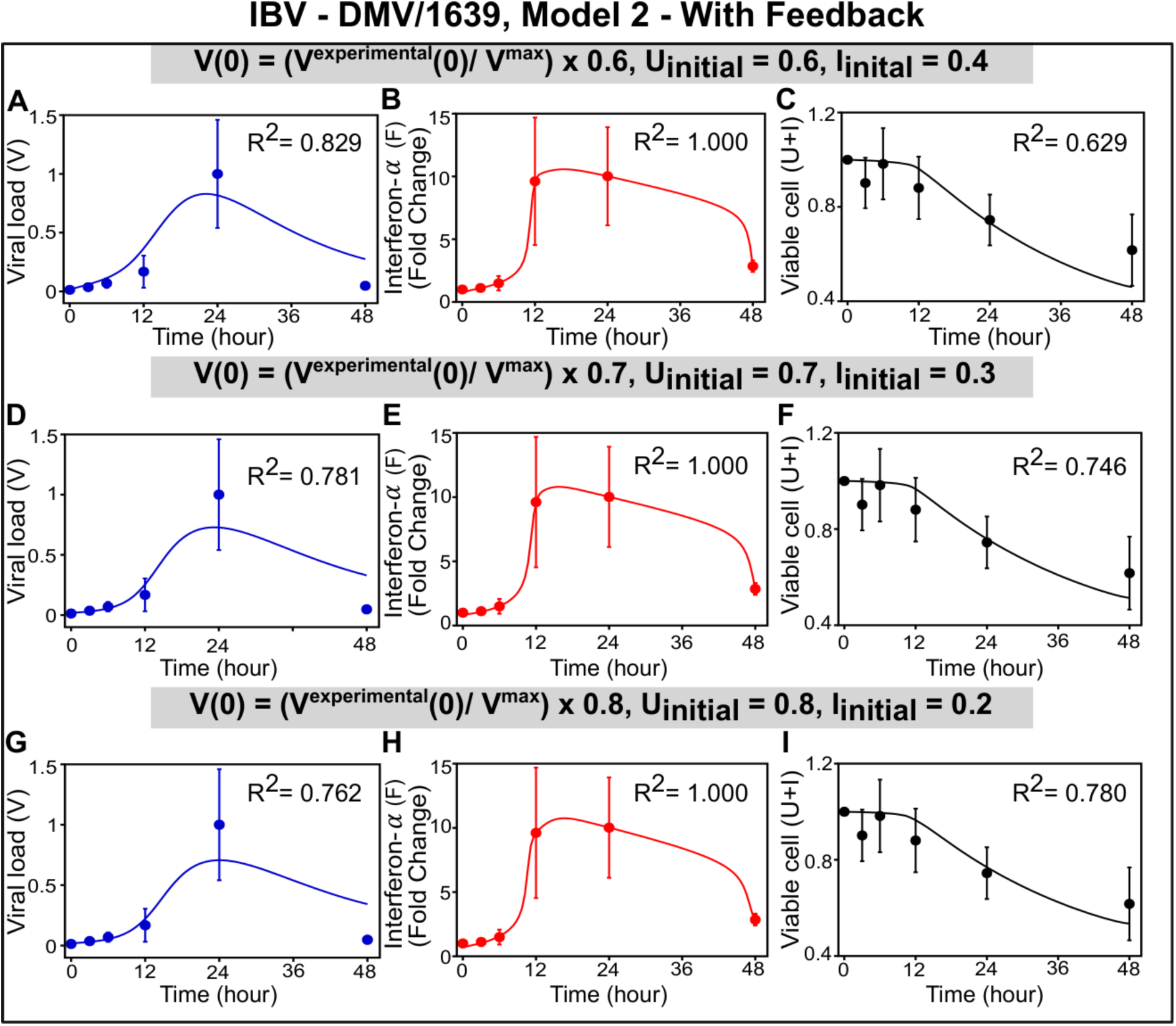
Influence of varying the fraction of initial extracellular virus and the fraction of uninfected and infected cells in Model 2 (with feedback) for the IBV-DMV/1639 strain. Predicted dynamics of extracellular viral load, interferon-α, and viable cells are shown by varying the fraction of the initial extracellular viral particles that can infect the cells. Note that the initial fraction of uninfected cells was taken to be the same as the initial extracellular virus fraction. These fractions were taken as 0.6 (A-C), 0.7 (D-F), and 0.8 (G-I), while the main figures were shown for a fraction of 0.5 --- see Equation S1 in the SI for the initialization of the Model. Solid lines represent model predictions, and filled circles are experimental data (error bars indicate standard deviation).

**Figure S6.**
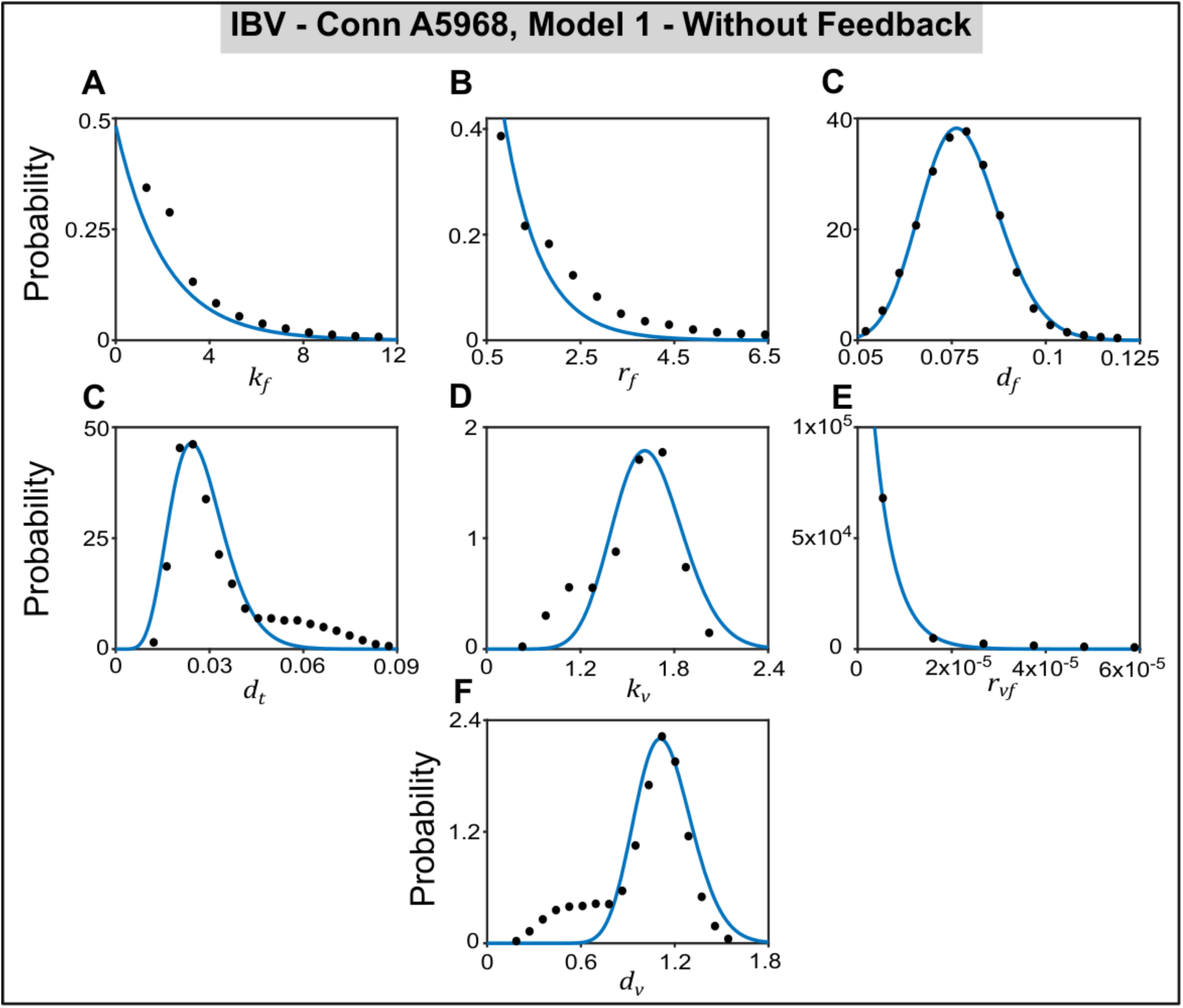
Distributions of the estimated Model 1 parameters for the IBV-Conn A5968 strain: Parameter distributions were obtained by repeatedly fitting the model to bootstrap-resampled experimental datasets. The resulting distributions were fitted with Gamma distributions (blue curves) to characterize the uncertainty in parameter estimation and were used to calculate the 95% confidence intervals shown in Table 1 (main text). The estimated parameter distributions were obtained for an initial extracellular virus fraction of 0.5 of the total initial viral load (as in Fig. 1, main text).

**Figure S7.**
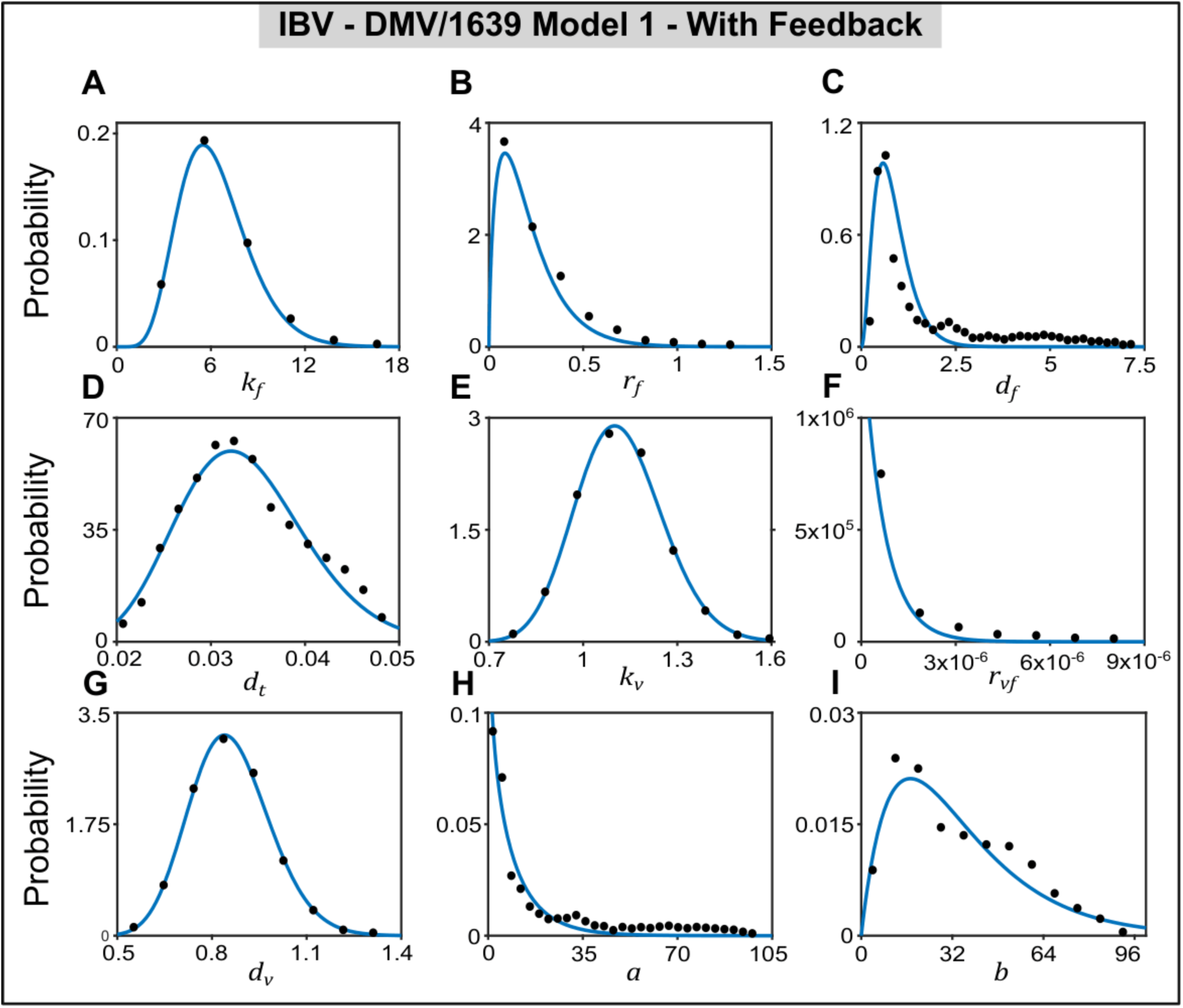
Distributions of the estimated Model 1 parameters (with feedback) for the IBV/DMV strain: Parameter distributions were obtained by repeatedly fitting the model to bootstrap-resampled experimental datasets. The resulting distributions were fitted with Gamma distributions (blue curves) to characterize the uncertainty in parameter estimation and were used to calculate the 95% confidence intervals shown in Table 1 (main text). The estimated parameter distributions were obtained for an initial extracellular virus fraction of 0.5 of the total initial viral load (as in Fig. 2, main text).

**Figure S8.**
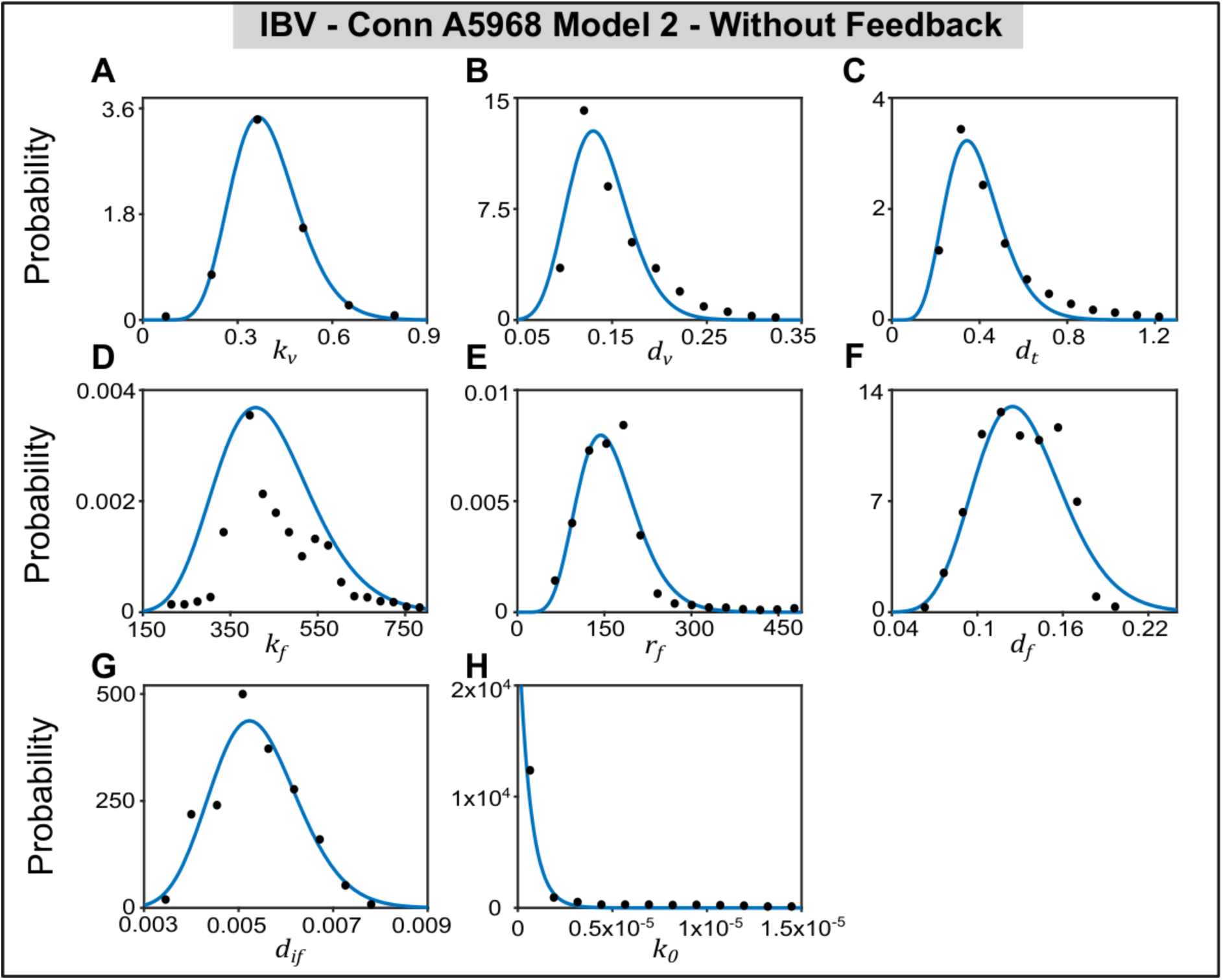
Distributions of the estimated Model 2 parameters for the IBV-Conn A5968 strain: Parameter distributions were obtained by repeatedly fitting the model to bootstrap-resampled experimental datasets. The resulting distributions were fitted with Gamma distributions (blue curves) to characterize the uncertainty in parameter estimation and were used to calculate the 95% confidence intervals shown in Table 2 (main text). The estimated parameter distributions were obtained for an initial extracellular virus fraction of 0.5 of the total initial viral load, while the initial fractions of uninfected and infected cells were each fixed at 0.5 throughout the analysis. (as in Fig. 3, main text).

**Figure S9.**
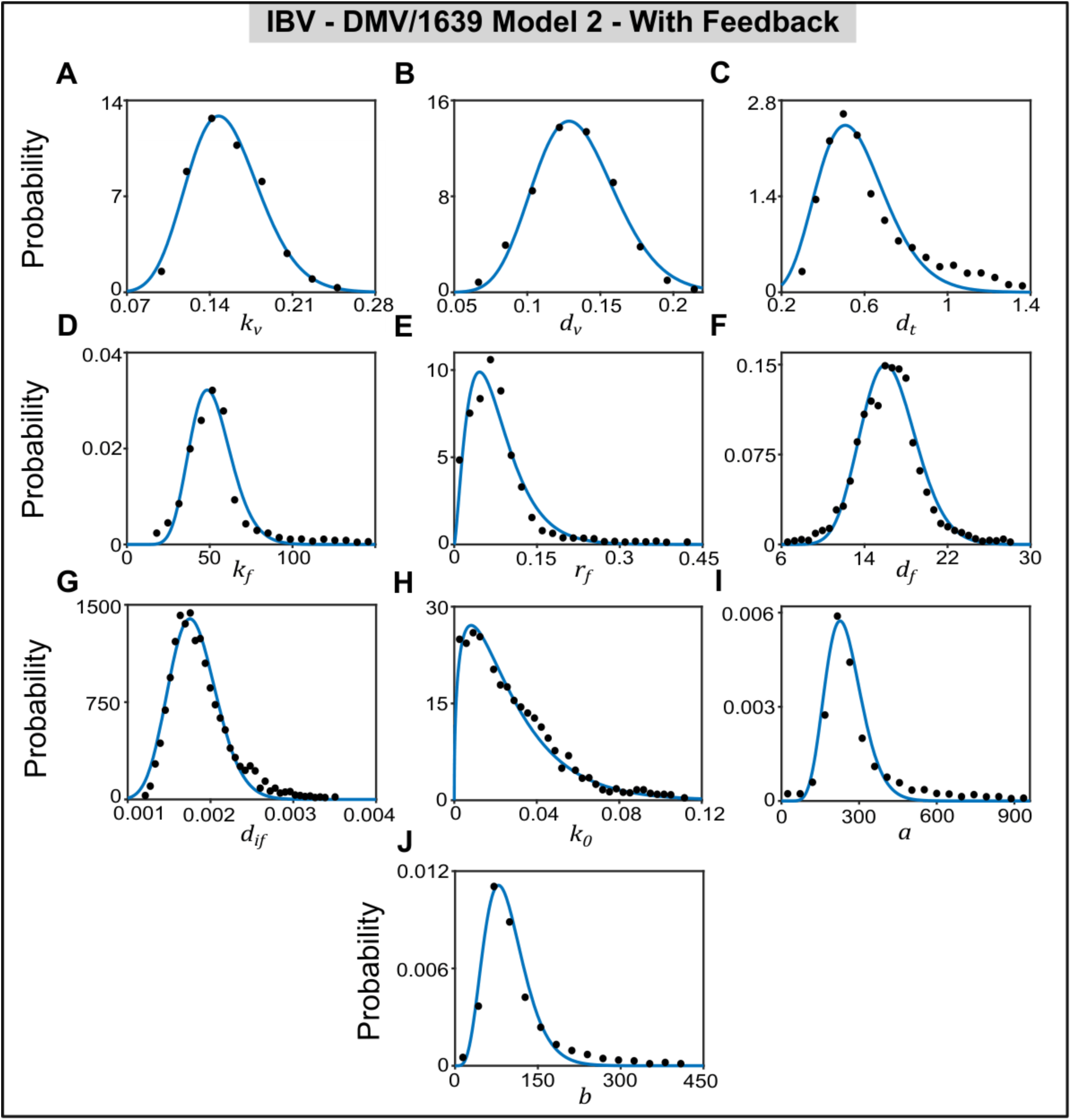
Distributions of the estimated Model 2 (with feedback) parameters for the IBV-DMV/1639 strain: Parameter distributions were obtained by repeatedly fitting the model to bootstrap-resampled experimental datasets. The resulting distributions were fitted with Gamma distributions (blue curves) to characterize the uncertainty in parameter estimation and were used to calculate the 95% confidence intervals shown in Table 2 (main text). The estimated parameter distributions were obtained for an initial extracellular virus fraction of 0.5 of the total initial viral load, while the initial fractions of uninfected and infected cells were each fixed at 0.5 throughout the analysis. (as in Fig. 4, main text).

**Table S1:**
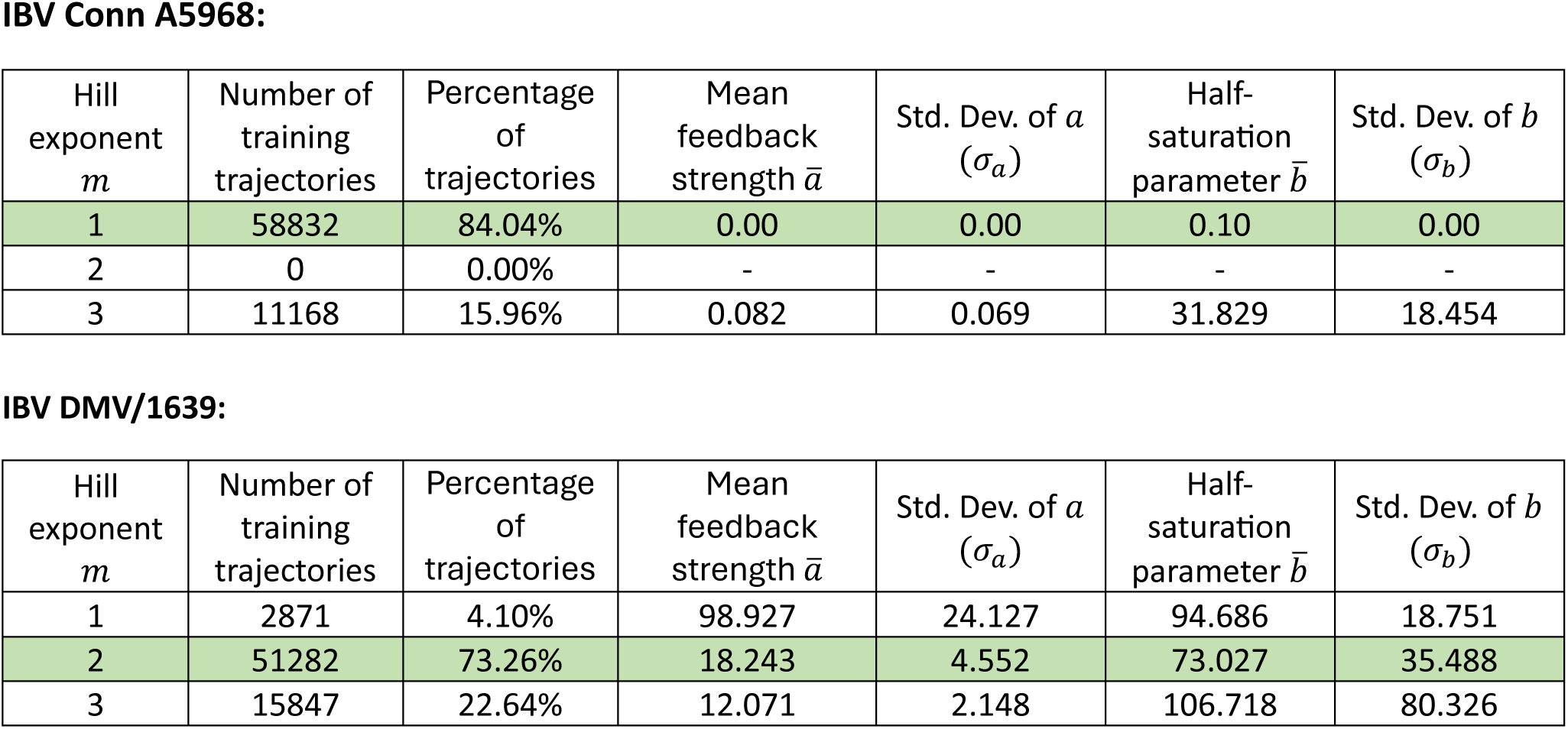
Mean values and standard deviations of the inferred feedback parameters across bootstrap trajectories.

| Hill exponent *m* | Number of training trajectories | Percentage of trajectories | Mean feedback strength *ā* | Std. Dev. of *a* (*σ*<sub>*a*</sub>) | Half-saturation parameter *b̄* | Std. Dev. of *b* (*σ*<sub>*b*</sub>) |
|---|---|---|---|---|---|---|
| **IBV Conn A5968:** |  |  |  |  |  |  |
| 1 | 58832 | 84.04% | 0.00 | 0.00 | 0.10 | 0.00 |
| 2 | 0 | 0.00% | - | - | - | - |
| 3 | 11168 | 15.96% | 0.082 | 0.069 | 31.829 | 18.454 |
| **IBV DMV/1639:** |  |  |  |  |  |  |
| Hill exponent *m* | Number of training trajectories | Percentage of trajectories | Mean feedback strength *ā* | Std. Dev. of *a* (*σ*<sub>*a*</sub>) | Half-saturation parameter *b̄* | Std. Dev. of *b* (*σ*<sub>*b*</sub>) |
| 1 | 2871 | 4.10% | 98.927 | 24.127 | 94.686 | 18.751 |
| 2 | 51282 | 73.26% | 18.243 | 4.552 | 73.027 | 35.488 |
| 3 | 15847 | 22.64% | 12.071 | 2.148 | 106.718 | 80.326 |

## Notes

### Competing Interest Statement

The authors have declared no competing interest.

## References

1. P. Baccam, C. Beauchemin, C. A. Macken, F. G. Hayden, A. S. Perelson, Kinetics of Influenza A Virus Infection in Humans. J. Virol. 80, 7590–7599 (2006).

2. J. H. Darbyshire, J. K. A. Cook, R. W. Peters, Comparative growth kinetic studies on avian infectious bronchitis virus in different systems. J. Comp. Pathol. 85, 623–630 (1975).

3. C. Hadjichrysanthou, et al., Understanding the within-host dynamics of influenza A virus: from theory to clinical implications. J. R. Soc. Interface 13, 20160289 (2016).

4. N. K. Vaidya, A. Bloomquist, A. S. Perelson, Modeling Within-Host Dynamics of SARS-CoV-2 Infection: A Case Study in Ferrets. Viruses 13, 1635 (2021).

5. J. K. Mirgichan, C. G. Ngari, S. Karanja, R. Muriungi, Mathematical modeling and simulation of hepatitis B transmission dynamics with passive immunity and control strategies. Heliyon 11, e41744 (2025).

6. K. A. Pawelek, et al., Modeling Within-Host Dynamics of Influenza Virus Infection Including Immune Responses. PLoS Comput. Biol. 8, e1002588 (2012).

7. P. Nagaraja, et al., The within-host fitness of HIV-1 increases with age in ART-naïve HIV-1 subtype C infected children. Sci. Rep. 11, 2990 (2021).

8. A. U. Neumann, et al., Hepatitis C Viral Dynamics in Vivo and the Antiviral Efficacy of Interferon-α Therapy. Science 282, 103–107 (1998).

9. B. O. Emerenini, S. C. Inyama, Mathematical model and analysis of hepatitis B virus transmission dynamics. F1000Research 7, 1312 (2018).

10. K. Best, D. H. Barouch, J. Guedj, R. M. Ribeiro, A. S. Perelson, Zika virus dynamics: Effects of inoculum dose, the innate immune response and viral interference. PLOS Comput. Biol. 17, e1008564 (2021).

11. S. Wang, et al., Modeling the viral dynamics of SARS-CoV-2 infection. Math. Biosci. 328, 108438 (2020).

12. J. Yang, et al., Parameter identifiability of a within-host SARS-CoV-2 epidemic model. Infect. Dis. Model. 9, 975–994 (2024).

13. I. M. Elbaz, H. El-Metwally, M. A. Sohaly, Viral kinetics, stability and sensitivity analysis of the within-host COVID-19 model. Sci. Rep. 13, 11675 (2023).

14. E. A. Hernandez-Vargas, J. X. Velasco-Hernandez, In-host Mathematical Modelling of COVID-19 in Humans. Annu. Rev. Control 50, 448–456 (2020).

15. C. A. Beauchemin, A. Handel, A review of mathematical models of influenza A infections within a host or cell culture: lessons learned and challenges ahead. BMC Public Health 11, S7 (2011).

16. R. Callard, A. J. T. George, J. Stark, Cytokines, Chaos, and Complexity. Immunity 11, 507–513 (1999).

17. K. Best, A. S. Perelson, Mathematical modeling of within-host Zika virus dynamics. Immunol. Rev. 285, 81–96 (2018).

18. R. Sachak-Patwa, E. I. Lafferty, C. J. Schmit, R. N. Thompson, H. M. Byrne, A target-cell limited model can reproduce influenza infection dynamics in hosts with differing immune responses. J. Theor. Biol. 567, 111491 (2023).

19. J. J. A. Weaver, J. E. Shoemaker, Mathematical Modeling of RNA Virus Sensing Pathways Reveals Paracrine Signaling as the Primary Factor Regulating Excessive Cytokine Production. Processes 8, 719 (2020).

20. L. G. Guidotti, F. V. Chisari, Cytokine-Mediated Control of Viral Infections. Virology 273, 221–227 (2000).

21. C. Leon, A. Tokarev, A. Bouchnita, V. Volpert, Modelling of the Innate and Adaptive Immune Response to SARS Viral Infection, Cytokine Storm and Vaccination. Vaccines 11, 127 (2023).

22. A. Handel, I. M. Longini, R. Antia, Towards a quantitative understanding of the within-host dynamics of influenza A infections. J. R. Soc. Interface 7, 35–47 (2010).

23. M. Sato, et al., Positive feedback regulation of type I *IFN* genes by the IFN-inducible transcription factor IRF-7. FEBS Lett. 441, 106–110 (1998).

24. Y. Wang, et al., Chicken interferon regulatory factor 7 (IRF7) can control ALV-J virus infection by triggering type I interferon production through affecting genes related with innate immune signaling pathway. Dev. Comp. Immunol. 119, 104026 (2021).

25. I. Kareva, F. Berezovskaya, G. Karev, Mathematical model of a cytokine storm. [Preprint] (2022). Available at: http://biorxiv.org/lookup/doi/10.1101/2022.02.15.480585 [Accessed 14 August 2025].

26. S. M. Suhail, et al., Regulatory T cell infiltration precedes early events associated with persistent infectious bronchitis virus (IBV) infection in the cecal tonsils of chickens. Virology 612, 110691 (2025).

27. K. O’Dowd, et al., Characterization of microRNA candidates at the primary site of infectious bronchitis virus infection: A comparative study of in vitro and in vivo avian models. PLOS ONE 20, e0319153 (2025).

28. M. Farooq, A. Ali, M. S. H. Hassan, M. F. Abdul-Careem, Nucleotide and Amino Acid Analyses of Unique Infectious Bronchitis Virus (IBV) Variants from Canadian Poultry Flocks with Drop in Egg Production. Genes 15, 1480 (2024).

29. F. Bande, et al., Pathogenesis and Diagnostic Approaches of Avian Infectious Bronchitis. Adv. Virol. 2016, 1–11 (2016).

30. M. S. H. Hassan, et al., Delmarva (DMV/1639) Infectious Bronchitis Virus (IBV) Variants Isolated in Eastern Canada Show Evidence of Recombination. Viruses 11, 1054 (2019).

31. X.-T. Xie, et al., A within-host mathematical model of H9N2 avian influenza infection and type-I interferon response pathways in chickens. J. Theor. Biol. 499, 110320 (2020).

32. L. Bordes, et al., Mathematical modeling of *in vitro* replication dynamics for multiple highly pathogenic avian influenza clade 2.3.4.4 viruses in chicken and duck cells. Microbiol. Spectr. 14, e00843-26 (2026).

33. S. Satani, H. M. Dobrovolny, Modeling the interferon response to avian influenza. Sci. Rep. (2026). 10.1038/s41598-026-71730-9.

34. R. Howey, et al., Modelling the within-host dynamics of the foot-and-mouth disease virus in cattle. Epidemics 4, 93–103 (2012).

35. C. E. Brook, et al., Accelerated viral dynamics in bat cell lines, with implications for zoonotic emergence. eLife 9, e48401 (2020).

36. O. Tomori, D. O. Oluwayelu, Domestic Animals as Potential Reservoirs of Zoonotic Viral Diseases. Annu. Rev. Anim. Biosci. 11, 33–55 (2023).

37. A. Ul-Rahman, H. M. Ishaq, M. A. Raza, M. Z. Shabbir, Zoonotic potential of Newcastle disease virus: Old and novel perspectives related to public health. Rev. Med. Virol. 32, e2246 (2022).

38. J. Baumann, N. M. Kouassi, E. Foni, H.-D. Klenk, M. Matrosovich, H1N1 Swine Influenza Viruses Differ from Avian Precursors by a Higher pH Optimum of Membrane Fusion. J. Virol. 90, 1569–1577 (2016).

39. M. S. H. Hassan, et al., Delmarva (DMV/1639) Infectious Bronchitis Virus (IBV) Variants Isolated in Eastern Canada Show Evidence of Recombination. Viruses 11, 1054 (2019).

40. M. E. Mahmoud, et al., Cyclooxygenase-2/Prostaglandin E2 Pathway Facilitates Infectious Bronchitis Virus-Induced Necroptosis in Chicken Macrophages, a Caspase-Independent Cell Death. Viruses 17, 503 (2025).

41. J. J. Sedmak, S. E. Grossberg, Interferon Bioassay: Reduction in Yield of Myxovirus Neuraminidases. J. Gen. Virol. 21, 1–7 (1973).

42. D. A. Messenger, D. M. Bortz, Weak SINDy: Galerkin-Based Data-Driven Model Selection. Multiscale Model. Simul. 19, 1474–1497 (2021).

43. S. L. Brunton, J. L. Proctor, J. N. Kutz, Discovering governing equations from data by sparse identification of nonlinear dynamical systems. Proc. Natl. Acad. Sci. 113, 3932–3937 (2016).

44. T. Cover, P. Hart, Nearest neighbor pattern classification. IEEE Trans. Inf. Theory 13, 21–27 (1967).

45. A. S. Perelson, R. Ke, Mechanistic Modeling of SARS-CoV-2 and Other Infectious Diseases and the Effects of Therapeutics. Clin. Pharmacol. Ther. 109, 829–840 (2021).

46. J. B. Xavier, et al., Mathematical models to study the biology of pathogens and the infectious diseases they cause. iScience 25, 104079 (2022).

47. L. Ungrová, et al., Avian interferon regulatory factor (IRF) family reunion: IRF3 and IRF9 found. BMC Biol. 23, 180 (2025).

48. Y. Cheng, et al., IRF7 Is Involved in Both STING and MAVS Mediating IFN-β Signaling in IRF3-Lacking Chickens. J. Immunol. 203, 1930–1942 (2019).

49. J. Kint, et al., Activation of the Chicken Type I Interferon Response by Infectious Bronchitis Coronavirus. J. Virol. 89, 1156–1167 (2015).

50. D. Das, D. Das, A. Prasad, Giant number fluctuations in microbial ecologies. J. Theor. Biol. 308, 96–104 (2012).

51. A. W. C. Yan, P. Cao, J. M. McCaw, On the extinction probability in models of within-host infection: the role of latency and immunity. J. Math. Biol. 73, 787–813 (2016).

52. F. S. Heldt, S. Y. Kupke, S. Dorl, U. Reichl, T. Frensing, Single-cell analysis and stochastic modelling unveil large cell-to-cell variability in influenza A virus infection. Nat. Commun. 6, 8938 (2015).

1. A. Amarasinghe, et al., Infectious bronchitis corona virus establishes productive infection in avian macrophages interfering with selected antimicrobial functions. PLOS ONE 12, e0181801 (2017).

2. M. E. Mahmoud, et al., Cyclooxygenase-2/prostaglandin E2 pathway regulates infectious bronchitis virus replication in avian macrophages. J. Gen. Virol. 105 (2024).

3. A. M. Kameka, S. Haddadi, D. S. Kim, S. C. Cork, M. F. Abdul-Careem, Induction of innate immune response following infectious bronchitis corona virus infection in the respiratory tract of chickens. Virology 450–451, 114–121 (2014).

4. D. A. Messenger, D. M. Bortz, Weak SINDy: Galerkin-Based Data-Driven Model Selection. Multiscale Model. Simul. 19, 1474–1497 (2021).

5. T. Cover, P. Hart, Nearest neighbor pattern classification. IEEE Trans. Inf. Theory 13, 21–27 (1967).

